# Revealing the unexpected interplay between the Proteasome Activator PA200 and the immunoproteasome

**DOI:** 10.1101/2025.03.18.643936

**Authors:** Dušan Živković, Fatme Mourtada, Angelique Sanchez Dafun, Ayse Seda Yazgili, Marijke Jansma, Przemysław Grygier, Michał Rawski, Anna Czarna, Stefan Bohn, Krzysztof M. Zak, Norbert Reiling, Linda Zemke, Kai Guo, Jürgen Behr, Odile Burlet-Schiltz, Grzegorz M. Popowicz, Julien Marcoux, Silke Meiners, Marie-Pierre Bousquet

**Affiliations:** Institut de Pharmacologie et de Biologie Structurale (IPBS), Université de Toulouse, CNRS, Université Toulouse (UT), Toulouse, France; Infrastructure Nationale de Protéomique, ProFI, UAR 2048, Toulouse, France; Research Center Borstel/Leibniz Lung Center, RG Immunology and Cell Biology, Airway Research Center North (ARCN), Member of the German Center for Lung Research (DZL), Parkallee 1-40, 23845, Borstel, Germany; Comprehensive Pneumology Center (CPC), University Hospital of the Ludwig -Maximilians-University (LMU) and Helmholtz Center Munich, Member of the German Center for Lung Research (DZL), Max-Lebsche Platz 31, 81377, Munich, Germany; Molecular Targets and Therapeutics Center, Institute of Structural Biology, Helmholtz Munich, Neuherberg, Germany; TUM School of Natural Sciences, Department of Bioscience, Bayerisches NMR Zentrum, Technical University of Munich, Garching, Germany; Malopolska Centre of Biotechnology, Jagiellonian University, Gronostajowa 7a, Krakow 30-387, Poland; Doctoral School of Exact and Natural Sciences, Jagiellonian University, Lojasiewicza 11, Krakow 30-348, Poland; National Synchrotron Radiation Centre SOLARIS, Jagiellonian University, Kraków, Poland; Cryo-Electron Microscopy Platform and Institute of Structural Biology, Helmholtz Munich; Ingolstädter Landstraße 1, 85764 Neuherberg, Germany; Research Center Borstel/Leibniz Lung Center, RG Microbial Interface Biology, Parkallee 1-40, 23845, Borstel, Germany, German Center for Infection Research (DZIF), Partner Site Hamburg-Lübeck-Borstel-Riems, 23845 Borstel, Germany; Department of Medicine V, LMU University Hospital, LMU Munich, Member of the German Center for Lung Research (DZL), Marchioninistr. 15, 81377, Munich, Germany; Institute of Experimental Medicine, Christian-Albrechts University Kiel, Kiel, Germany

## Abstract

The proteasome activator PA200 binds to the catalytic core of the proteasome, the 20S, and activates its proteolytic activities. The cellular function of PA200 is poorly understood and appears to be cell type and differentiation specific. Recent evidence suggests that PA200 not only binds to the standard 20S (s20S) proteasome but also to the specialized immunoproteasome (i20S) which plays a key role in anti-viral and anti-tumor immunity.

We here investigated the interaction of PA200 and the immunoproteasome in detail. We show the very first cryo-EM structures of the singly- and doubly-capped i20S-PA200 complexes that revealed no major difference regarding the first binding event of PA200 to the i20S vs. the s20S. However, first PA200 binding triggered a subtle and long range allosteric bending of the i20S barrel which was not seen in the s20S-PA200 complexes. This resulted in major structural rearrangements in the opposite unbound α ring - the displacement of atoms up to 5.4 Å and the increase in its outer diameter - thereby increasing the occupancy of the second PA200 binding site. Mass photometry confirmed higher occupancy of PA200 to the i20S versus the s20S. Binding of PA200 to the i20S enhanced proteasomal activation compared to the s20S. Co-expression of PA200 and the i20S in cells and tissues, however, is restricted but their interaction is favored upon co-expression. The expression of PA200 and the catalytic subunits of the i20S is differentially regulated depending on the cellular context. Our data also suggest that PA200 has the potential to regulate i20S gene expression whereas the i20S has no effects on PA200 expression. Overall, this work sheds new light on the interaction of PA200 with the i20S from a structural, mechanistic and cellular point of view. Importantly, we identify PA200 as a key regulator of the i20S whenever PA200 and the catalytic subunits of the i20S are co-expressed in the same cell.

## Introduction

The proteasome is the main protease for targeted protein degradation in the cell. It degrades old, un-used and damaged proteins into 8-21 amino acid long peptides. These are further cleaved into amino acids for amino acid recycling by the ribosome but are also used as antigens for presentation on major histocompatibility complex (MHC) class I molecules to the immune system, i.e. CD8 T cells^1^. The proteasome is composed of the catalytic core, the 20S proteasome, and associated proteasome regulators^2^. As such, the term proteasome refers to a family of complexes which can rapidly assemble and thereby adjust substrate specificity, proteolytic activities and subcellular localization of proteasome-dependent protein degradation according to cellular needs^3^.

The 20S proteasome is a barrel-shaped protease composed of two outer α and two inner β rings of seven subunits each^4^. Three of the seven β subunits form the proteolytically active sites which contain N-terminal threonine residues as active centers. These β1, β2, and β5 subunits of the standard 20S (s20S) proteasome possess caspase-like (C-L), trypsin-like (T-L), and chymotrypsin-like (CT-L) activities cleaving proteins after acidic, basic and hydrophobic amino acids, respectively. An alternative set of catalytic subunits (β1i, β2i, β5i) can be incorporated into the 20S proteasome forming the immunoproteasome (i20S)^5^. These three immunosubunits are transcriptionally induced in cells upon viral infections or inflammatory signaling and possess an important role in immune responses including MHC class I-mediated antigen presentation^5^. Immune cells contain predominant amounts of i20S and varying amounts of s20S^6,7^.

Amongst the proteasome regulators, the 19S activator is most predominant in the cell and mediates ubiquitin- and ATP-dependent protein degradation^8,9^. It forms the 26S proteasome upon assembly with the 20S core particle. Other regulators include PA28αβ and PA28γ as well as PA200, also known as PSME4^5^. While the cellular function of PA28αβ in immune activation and of PA28γ in nuclear protein degradation are well studied^3^, the function of PA200 is still enigmatic^10^. PA200 can bind to the 20S catalytic core either on one side or on both sides forming singly- or doubly-capped 20S proteasome complexes, respectively^11,12^. It can also bind to the free end of the 26S proteasome complex^13^. While structural data clearly reveal a proteasome activating role of PA200, cellular data on which active sites are activated by PA200 are conflicting^11,14^. Moreover, the cellular function of PA200 appears to be cell type and differentiation specific, ranging from DNA damage repair and chromatin remodeling^15–17^ to multiple roles in aging^18^, differentiation^19^, mitochondrial and protein stress responses^13,20–22^. Together with the Merbl lab, we recently demonstrated that PA200 binds to the i20S and thereby modulates antigen diversity to abrogate antitumor immunity in lung cancer^14^.

We here investigated the interplay of PA200 and the immunoproteasome in detail by combining structural analysis of i20S-PA200 complexes with biochemical and cellular analyses. We provide the very first cryoEM structure of the i20S singly or doubly-capped with PA200 that exhibit striking structural differences compared to their s20S counterparts. These major structural rearrangements result in enhanced formation of doubly-capped i20S–PA200 complexes with increased activation of the i20S compared to the s20S upon binding of PA200 *in vitro*. Co-expression of PA200 and the i20S is restricted and regulation of the expression of PA200 and the catalytic subunits of the i20S depends on the cellular context. Our data also suggest that PA200 has the potential to regulate i20S gene expression and thereby 20S proteasome composition.

## Results

### Structural analysis of immunoproteasome-PA200 complexes

We obtained high resolution structures for both, singly- and doubly-capped i20S-PA200 complexes (at 2.85 Å and 2.89 Å resolution, respectively). The excellent quality of our data prompted us to solve the structures of both complexes and compare the i20S-PA200 interaction with structures of the s20S-PA200 complex, recently resolved by three groups^11,12,23^. The doubly-capped structure shows a high degree of similarity with s20S-PA200 complexes with no major differences at the interface between PA200 and 20S proteasomes (Figure 1A-B, see Figure S1 for details on 3D classification). However, our structure shows a remarkable “bend” of the entire complex. This is best seen when the structures are aligned over one PA200 molecule. With perfect alignment on one side of the doubly-capped complex, the other side of the complex reveals marked misalignment that enlarges with distance from the aligned molecule (Figure 1C). To our knowledge, this bend has not been observed in any proteasome structure before. This is remarkable as previously resolved structures show strong similarity even when solved with different experimental methods (NMR, X-ray diffraction, cryo-EM). Importantly, the same structural shifting/tilting behavior was evident in singly-capped i20S-PA200 structures (Figure 2A and S2A-C), implying that binding of the first PA200 molecule is sufficient to induce this long-range allosteric change that is not further affected by the second PA200 binding event. Allosteric rearrangement results in allosteric displacement of α-ring subunits at the opposing end of the i20S (Figure 2A) but is not visible in the s20S-PA200 complex (Figure 2B). For example, the Cα of glutamine residue GLN149 in the proteasome α3 subunit of the upper α-ring shifts by 1.6 Å when s20S-PA200 and i20S-PA200 complexes are compared. The same atom in the lower α-ring, however, shifts by 5.0 Å and the diameter of the opposite unbound α-ring is allosterically increased by up to 3.5 Å (with some atoms displaced by up to 5.4 Å). It is tempting to speculate that this widening of the unbound α-ring might alter the binding of subsequent regulatory particles. Proteasome activators typically bind to the α subunits of the 20S core to open the inner channel enabling substrate access. The i20S-PA200 complex shows a degree of channel opening comparable to that of the s20S-PA200 (28-32 Å; Figure 2C-D), which - as expected - is significantly higher than that of the apo s20S and i20S complexes, but also remarkably larger than that of the only partially open s20S-PA28αβ^23^ and i20S-PA28αβ^24^ complexes (9-10 Å). This indicates that PA200 opens the gate of the s20S and i20S particles to a wider extent than PA28αβ. Detailed analysis of the catalytic sites also revealed significant differences when comparing the structures of the i20S vs i20S-PA200 complexes (Figure 2E), confirming that activator binding induces long distance structural changes to the catalytic sites as seen before^11,25^.

**Figure 1:**
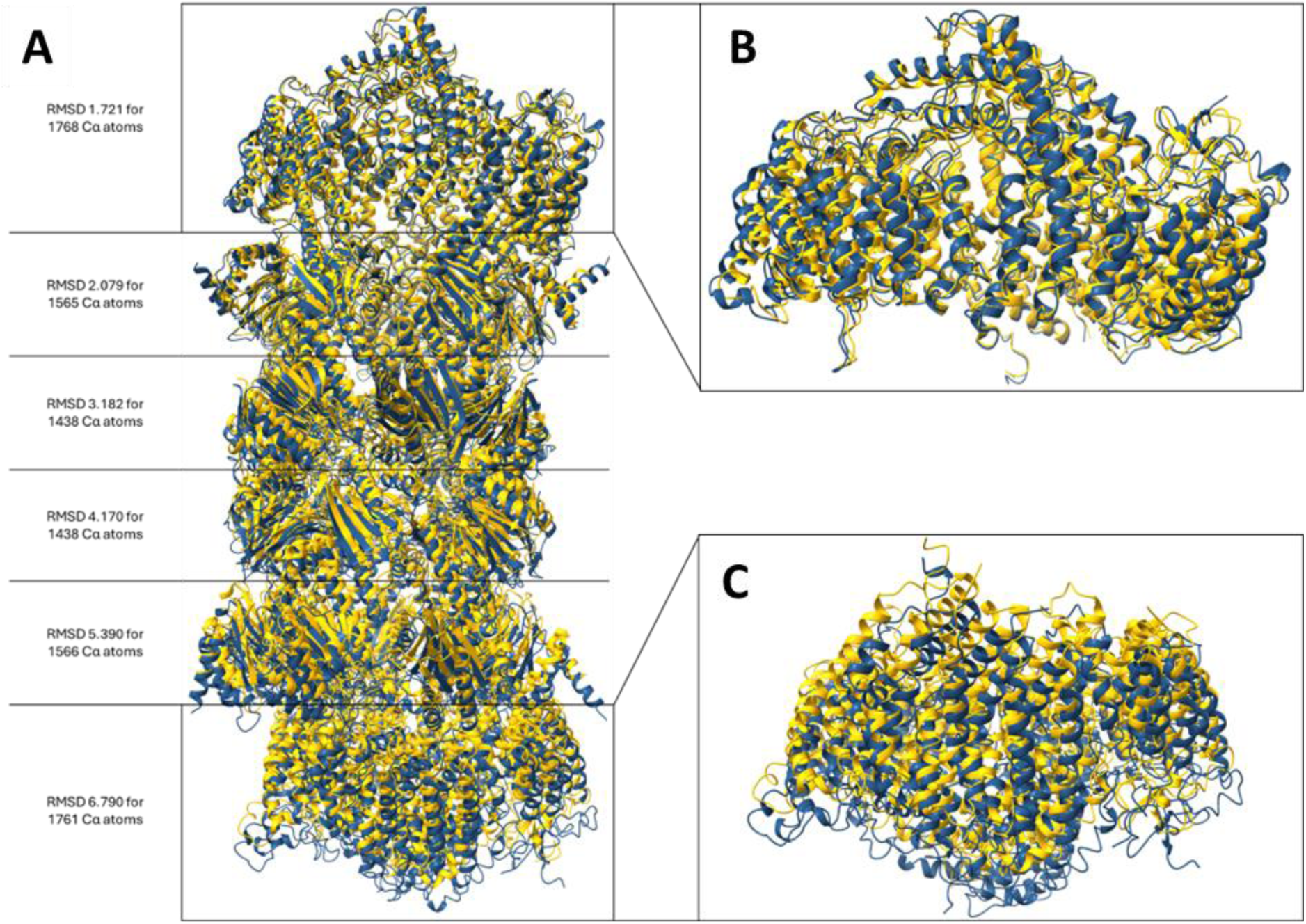
Cryo-EM analysis of the i20S-PA200 proteasome complex. The structure of doubly-capped i20S shares overall similarity to the s20S-PA200 structures reported before. (**A**) Overlay of the i20S-PA200 (blue) and s20S-PA200 (yellow, PDB: 6REY). (**B**) The structures were aligned by the PA200 molecule at the top. The overall axis of the complex is bent. (**C**) The subunits further away from aligned PA200 binding show distinct alterations.

**Figure 2:**
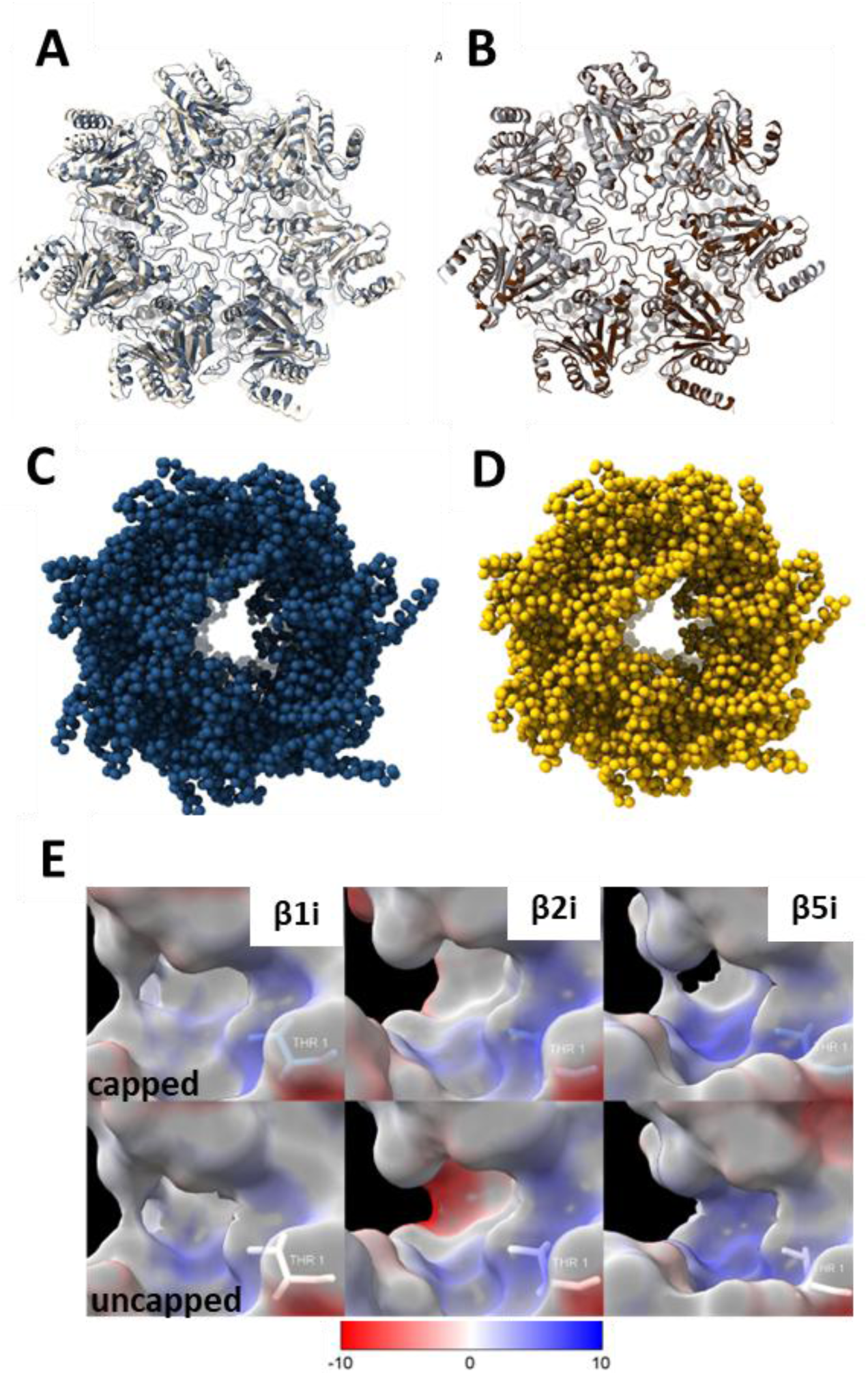
Binding of PA200 induces channel opening of i20S, allosterically alters the catalytic site and widens the opposite unbound α-ring diameter. Superimposition of the (**A**) unbound side of the singly-capped i20S-PA200 (white, PDB ID: XXXX) with uncapped i20S (blue, PDB ID: 6AVO) and (**B**) unbound side of the singly-capped s20S-PA200 (grey, PDB ID: 6KWY) with the uncapped s20S (brown, PDB ID: 6RGQ), (**C**) Structure of i20S-PA200 in an open state caused by PA200 binding. (**D**) s20S-PA200 in open state (PDB ID 6REY), (**E**) Comparison of active sites of the capped (top) and uncapped (bottom) i20S colored by electrostatics (positive in blue and negative in red).

Finally, we detected two molecules of inositol hexakisphosphates (IP6, phytic acid) binding to the positively charged grooves of PA200 (Figure S3), whereas previous structures of s20S-PA200 identified both IP6 and (5,6)-bisdiphosphoinositol tetrakisphosphate (5,6[PP]2-IP4)^11,12,23^.

### PA200 binds and activates the i20S more efficiently than the s20S

We further quantified the binding of PA200 to the i20S vs. s20S using mass photometry. This interferometric microscopy-based technology can measure molecular weights of single molecules (>40 kDa) in solution at nM concentrations, without any previous labeling^26,27^. For that, purified human i20S and s20S were incubated for 2 h with increasing amounts of recombinantly expressed and purified human PA200 at increasing PA200:20S molar ratios from 0 to 12.

We detected free 20S, singly-capped and doubly-capped proteasome complexes (Figure 3A). As expected, the total occupancy of the 20S increased with the PA200 to 20S ratio. We observed a higher occupancy of the i20S by PA200, compared to the s20S (Figure 3B). This titration allowed us to estimate the Kds of PA200 for the i20S and the s20S which were around 20-30 nM for the first binding event and 40-60 nM for the second binding event (Figure S4). We next determined the activation of i20S vs. s20S by PA200 using fluorogenic peptides for the three active sites. Activities were determined for *in vitro* complexes at increasing PA200:20S molar ratios (ranging from 0 to 8). As expected from our mass photometry data, binding of PA200 to the i20S resulted in the activation of the three main types of 20S proteolytic activities, namely the CT-L, T-L, and C-L. In particular, the PA200-bound i20S was significantly more active towards CT-L and T-L sites than the PA200-capped s20S at the highest molar ratios (Figure 4). Moreover, while the baseline CT-L and T-L activities of the i20S and s20S were similar, the i20S showed very low C-L activity compared to the s20S (Figure 4A), in line with published data^28^.

**Figure 3:**
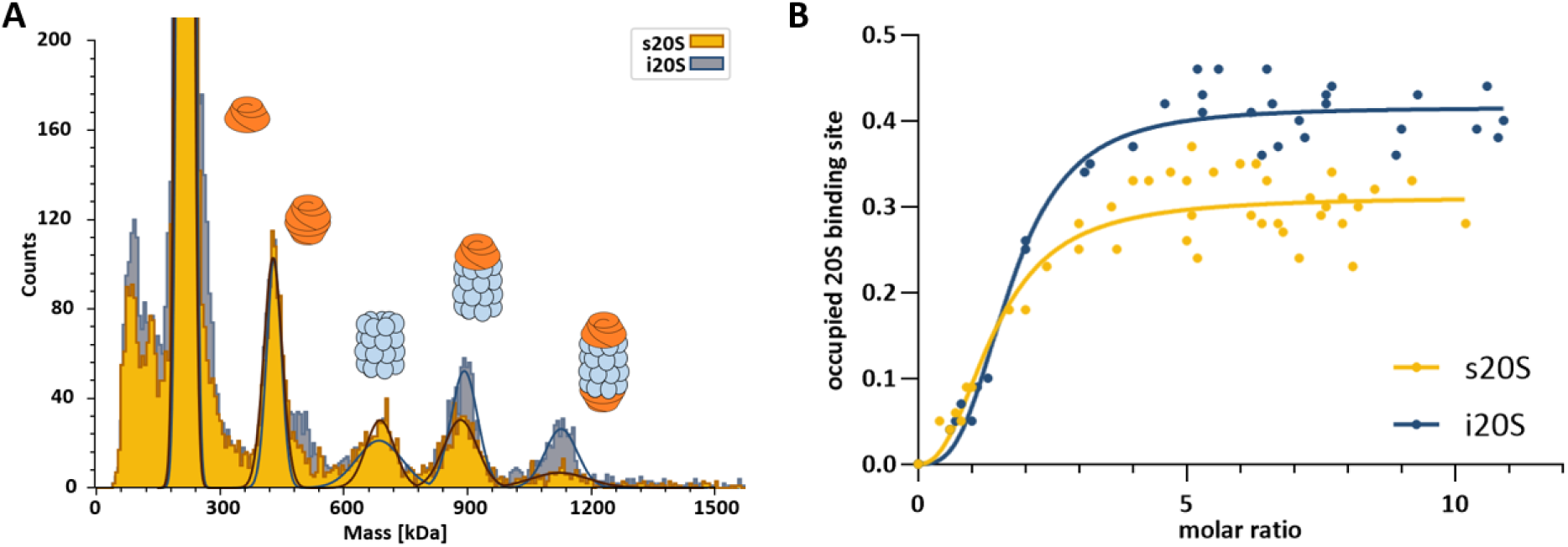
i20S-PA200 particles are preferentially formed compared to s20S-PA200 complexes. **(A)** Mass photometry allowed us to semi-quantify the relative abundances of free 20S (720 kDa), singly-capped 20S (930 kDa), and doubly-capped 20S (1,140 kDa). The used PA200:20S molar ratio was 8. **(B)** Titrations of PA200 to either s20S and i20S showed a higher total occupancy of the i20S. The total 20S occupied binding sites were calculated by summing the fractions of singly- and doubly-capped 20S (shown in Fig S3) and as detailed in the Materials and Methods section.

**Figure 4:**
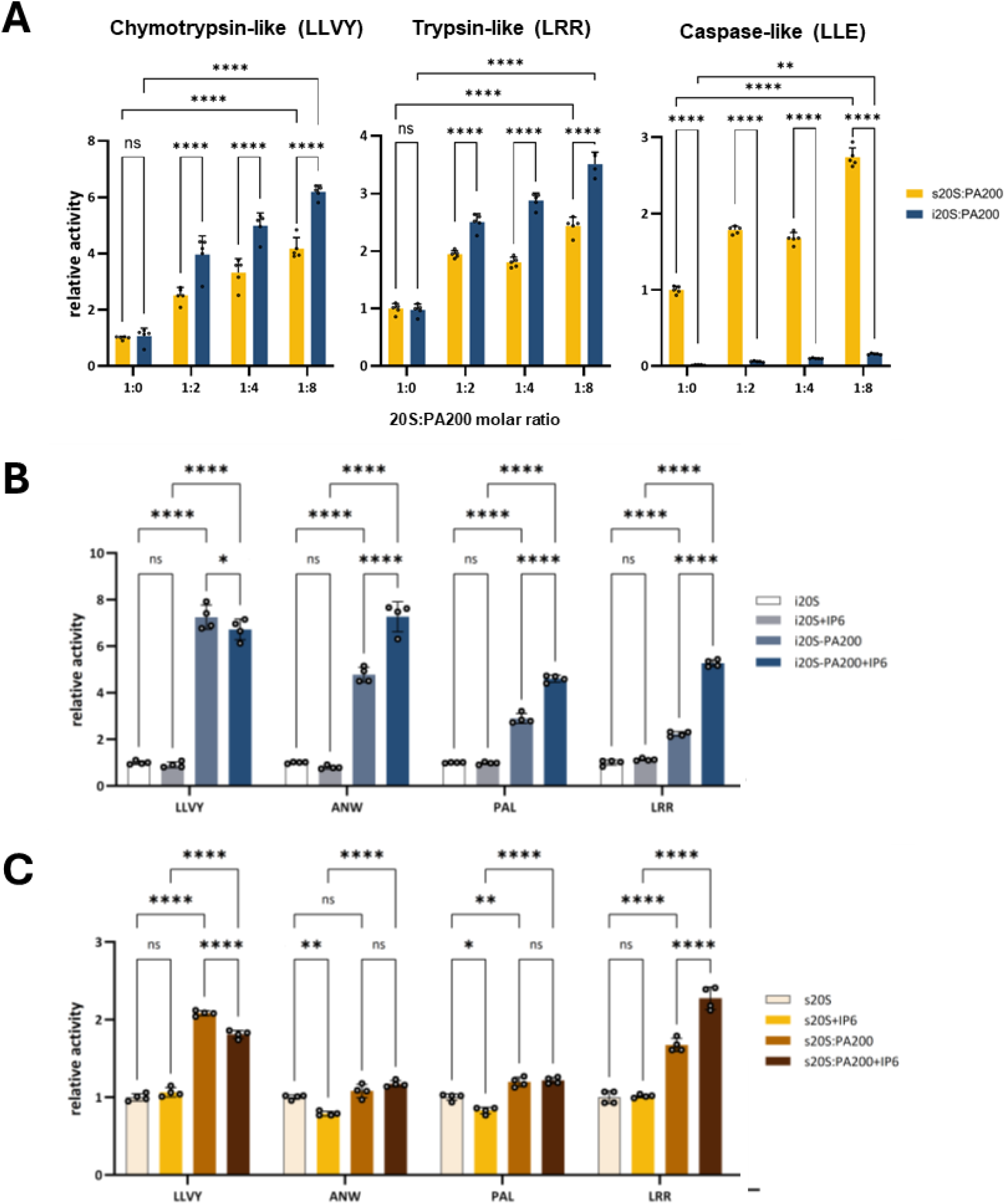
PA200 activates better the i20S and is further activated by phytic acid. (**A**) The three proteasome activities were assayed using isolated i20S and s20S incubated with increasing amounts of recombinantly expressed PA200 (molar ratios 1 to 8). The LLVY-AMC, LRR-AMC and LLE-AMC substrates were used to quantify the CT-L, T-L and C-L activities, respectively. The activity measured with the unactivated s20S was used as the reference activity. **(B-C)** Effect of phytic acid (IP6) on PA200-activated i20S (B) and s20S (C) catalytic activities using fluorogenic substrates specific for the distinct active sites of the i20S (LLVY, ANW, PAL, LRR substrates for**β**5i, **β**5i, **β**1i, **β**2i, respectively). PA200 was incubated first with IP6 and then with 20S at final concentrations of 21.5 nM, 70 μM and 1.5 nM for PA200, IP6 and 20S, respectively. The activity measured with the unactivated s20S was used as the reference activity. The activity assays were performed in at least 4 replicates. Two-way ANOVA with Tukey’s multiple comparison test at 95% confidence interval was used with ns: no significance, *: p<0.05, **: p<0.01, ***: p<0.001, and ****: p<0.0001.

It was suggested that the IP6 and 5,6[PP]2-IP4 identified in the s20S-PA200 structures could be involved in regulating the activity of PA200^10–12^. In order to test this hypothesis, we assessed the effect of phytic acid on the activity of the i20S and s20S in the presence and absence of PA200 using different fluorogenic substrates specific for the distinct active sites in an *in vitro* substrate assay^9,29^. Despite a slight but significant decrease of the CT-L activity (LLVY substrate) for both 20S subtypes, our results show a strong increase of the three activities that are specific of the i20S, namely β1i (PAL), β2i (LRR) and β5i (ANW), in the presence of IP6 (Figures 4B-C).

In summary, single or double capping of the i20S with PA200 not only affects the catalytic sites, as seen with the s20S, but also induces a massive induced fit change that was not observed in other 20S structures and that could influence binding of a second PA200 molecule to the opposite free α-ring, as well as specificity of the i20S indirectly. Furthermore, we show here that phytic acid could indeed be a cofactor of i20S-PA200, as it increases its activity *in vitro*.

Our structural and *in vitro* activity assays thus suggest that binding of the first PA200 induces an allosteric widening of the opposite unbound α-ring, resulting in a higher binding occupancy of the i20S compared to the s20S. This results in a more efficient and stronger activation of all three active sites compared to the s20S, and more specifically of the CT-L and T-L activities that favor the generation of MHC-I antigenic peptides^30,31^.

### Enriched binding of i20S to PA200 in tissues

In a next step, we aimed to confirm our *in vitro* data on the favored interaction of the i20S with PA200 in cells and tissues. As PA200 is highly expressed in testis, we decided to revisit our previously generated dataset from bovine testes (PXD027436)^29^ to study the interaction of PA200 with the i20S in tissues under physiological conditions. Our interactome analysis of immunoprecipitated PA200 from bovine testes demonstrated enriched incorporation of the i20S subunits PSMB8-10 (Figure 5A, in blue) compared to their s20S counterpart subunits PSMB5-7 (Figure 5A, in yellow) into PA200-containing proteasome complexes. We confirmed enriched assembly of i20S with PA200 by comparing our PA200-pulldown data with an interactome obtained upon immunoprecipitation with an anti-α2 antibody that binds to an α subunit of the 20S proteasome and thereby pulls down all proteasome complexes within the tissue^9,32^. As shown in Figure 5B, the ratios of the i20S/s20S catalytic subunits were significantly higher in the anti-PA200-coIP compared to the anti-α2-coIP samples.

**Figure 5:**
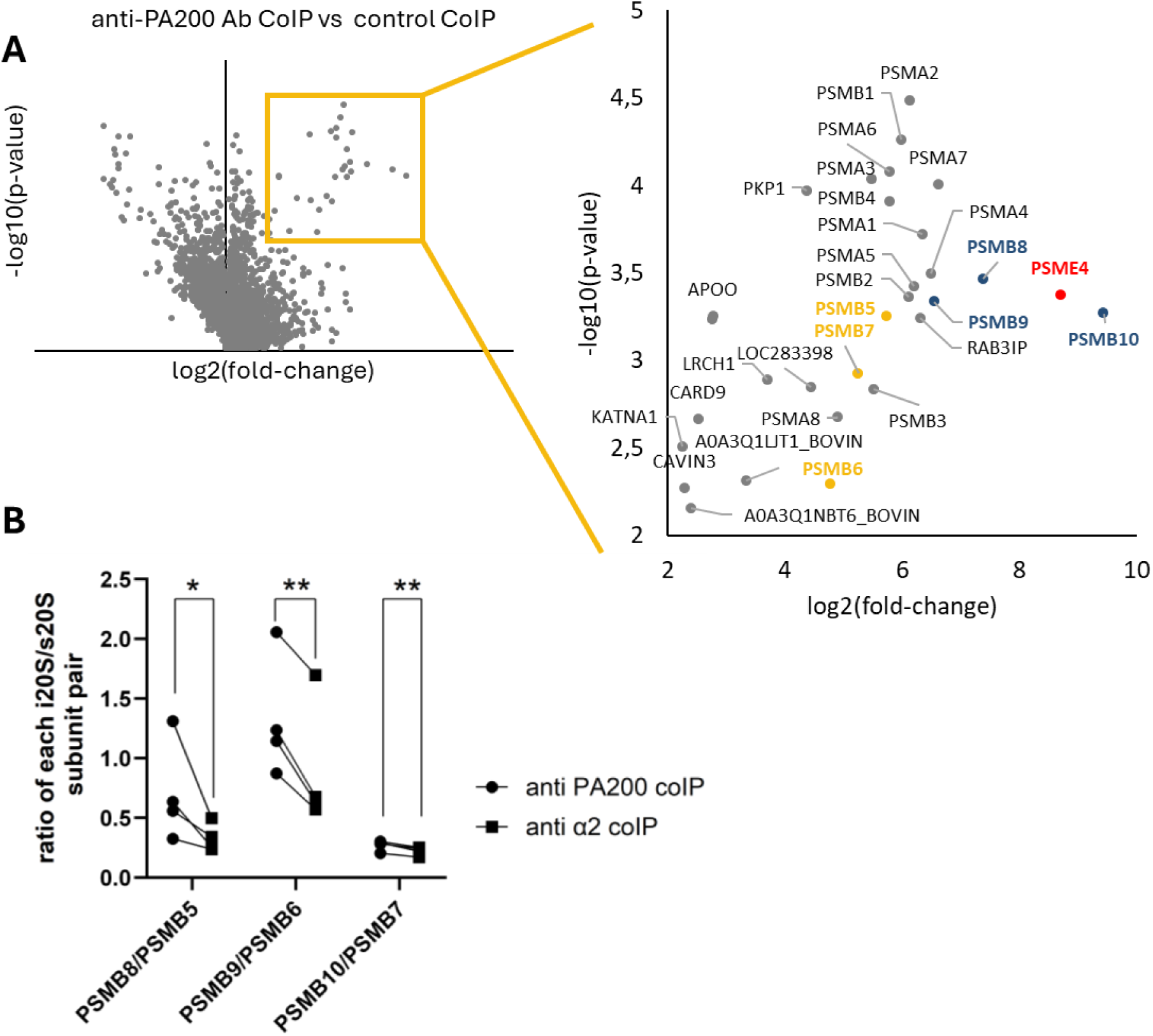
Enrichment of i20S in PA200-containing proteasome complexes in testis. **A)** Immunoprecipitation of PA200 in bovine testis and subsequent MS-based interactome analysis revealed enriched binding of the i20S catalytic subunits (β1i, β2i and β5i, ie PSMB8-10 respectively, labeled in yellow) compared to their s20S counterparts (β1, β2 and β5, ie PSMB5-7 respectively). The control co-IP was performed using the OX8 IgG1 unrelated monoclonal antibody directed against rat CD8 alpha, as published earlier (PXD027436)^29^ **B)** Quantification of the ratios of the corresponding catalytic i20S/s20S subunits upon PA200 pulldown of testes tissue and compared to total 20S pulldown of the same testes samples using an anti-α2 antibody. i20S subunits are significantly enriched compared to their s20S counterparts. A paired, two-sided t-test was used with *: p<0.05, **: p<0.01, n=4.

Given the fact that the anti-α2 antibody captures all 20S-containing proteasome complexes^32^, these data confirm that PA200 preferentially binds to the i20S compared to the s20S, which is fully in line with our *in vitro* data on structural interactions and activation presented above.

### Regulation of i20S-PA200 interaction on the cellular level

The key question is thus under which cellular conditions does PA200 bind to the i20S to regulate its activity. Under physiological conditions, the expression pattern of PA200 and the i20S hardly overlaps. According to the human proteome map^33^, PA200 (PSME4) is highly expressed in adult and fetal reproductive organs such as ovaries and testes, while the i20S subunits are abundantly expressed in immune cells such as monocytes, CD4, CD8, and natural killer (NK) cells (Figure S5).

Based on these data, we hypothesized that the interplay of PA200 and the immunoproteasome could be regulated at the cellular level via their expression. We thus tested two cellular model systems that were previously shown to regulate PA200 and i20S expression, respectively. To analyze the regulation of the i20S under conditions of PA200 induction, we used primary human lung fibroblasts (phLF) that were treated with TGF-β1 to upregulate PA200^19^. Vice versa, we investigated PA200 regulation under conditions of i20S activation in interferon gamma (IFNγ) stimulated phLF and upon infection of murine bone-marrow derived macrophages with *Mycobacterium tuberculosis* (*Mtb*)^34^.

On the RNA level, TGF-β1 treatment upregulated PA200 in several primary lung fibroblast (phLF) lines but uniformly downregulated the three i20S catalytic subunits PSMB8-10 (Figure 6A). Elevated incorporation of PA200 reduced assembly of i20S subunits which was confirmed by an interactome analysis of TGF-β1-treated phLFs compared to untreated controls (Figures 6B-C), using the anti-α2 antibody for pulldown of all crosslinked proteasome complexes. phLFs were also stimulated with IFNγ, resulting in the upregulation of the i20S, as indicated by a significant increase in PSMB8-10 expression, but without any impact on PSME4 mRNA levels as determined by RTqPCR (Figure 6D). Similarly, *Mtb* infected mouse macrophages strongly upregulated the i20S subunits PSMB8-10 but did not alter PSME4 RNA expression (Figure 6E). These data support the notion that PA200 and the i20S are differentially regulated under conditions of differentiation, cytokine treatment and infection.

**Figure 6.**
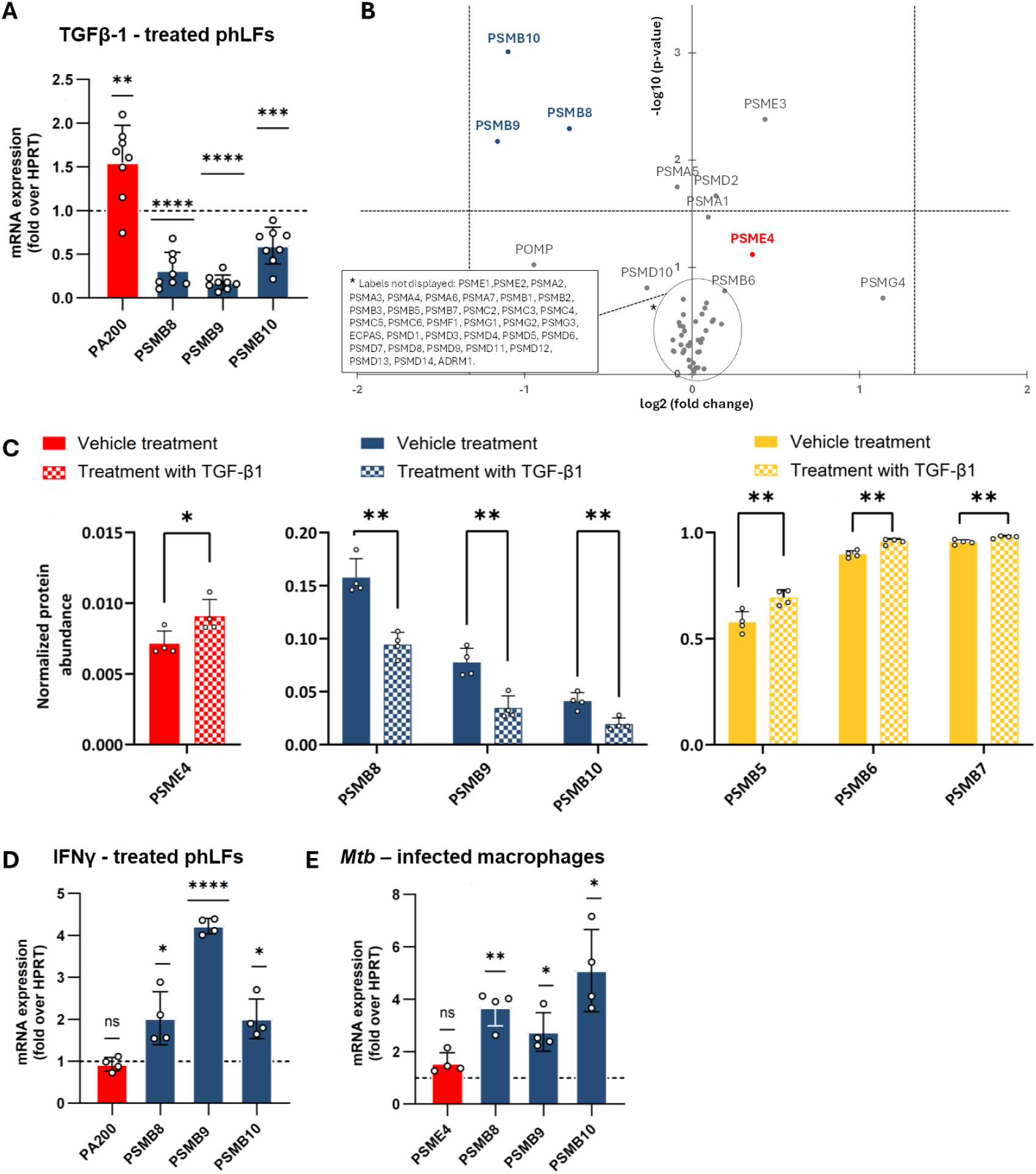
Interplay between PA200 and i20S expression in physiological contexts. RTqPCR-mediated quantification of PSME4 (PA200) and PSMB8-10 (β1i, β2i and β5i) in primary human lung fibroblasts (phLF) treated with **(A-C)** TGF-β1 or **(D)** IFNγ or in **(E)** mouse macrophages infected by *Mycobacterium tuberculosis Mtb*. phLFs were treated with TGF-β1 or IFNγ for 24 hours using several phLF cell lines derived from different donors (8 independent experiments). Murine bone-marrow derived macrophages were infected with *Mtb* for 24 h at a ratio of 3:1 (4 independent experiments) and RTqPCR analysis was performed. For **A**, **D** and **E**, expression of the target gene was normalized to Hypoxanthine phosphoribosyltransferase (HPRT) and the respective untreated control using the 2^-ΔΔCT^ method. The control was set to 1, as indicated with the dashed line. One sample t-test was applied with *: p<0.05, **: p<0.01, ***: p<0.001, ****: p<0.0001. **(B)** Volcano plot comparison of proteomic analyses of the proteasome co-immunoprecipitations (CoIPs) of phLFs treated with TGF-β1 (right) vs. vehicle-treated respective controls (left). Anti-α2 proteasome subunit antibody was used for the CoIPs. Paired t-test was used. Proteomic data was filtered to display only the proteasome complex subunits and proteasome-regulating proteins. Gene names PSMB8, PSMB9, PSMB10 and PSME4 are used instead of unique protein identifiers for ease of readability. **(C)** Variations of abundances of CoIPed PA200, immunosubunits and standard subunits upon TGF-β1 treatment, as determined by MS quantification (see Materials and Methods section).

### Regulation of PA200 by the immunoproteasome and vice versa

We finally addressed the question whether PA200 itself is involved in the regulation of the i20S and vice versa. For that we used our previously described PA200 knockout (KO) in two human lung cancer cell lines^27^. Genetic depletion of PA200 in A549 and H1299 reduced expression of the β1i catalytic subunit (PSMB9 gene) in both its pre-mature and mature forms (Figures 7A-B). These data were confirmed upon pulldown of 20S complexes using the anti-α2 antibody and interactome analysis demonstrating that the absence of PA200 impairs the assembly of β1i containing i20S complexes (Figure S6A). This resulted in an altered cellular composition of 20S complexes with reduced formation of intermediate proteasomes containing the two immunocatalytic subunits β1i and β5i (DIP, β1iβ5i i20S) and an increase in complexes harbouring only the immunocatalytic subunit β5i (SIP, β5i i20S) (Figure S6A)^35,36^. Concerted and even more pronounced downregulation of the i20S catalytic subunits PSMB8 and PSMB9 was observed on the RNA level upon transient silencing of PA200 in A549 cells (Figure S6B). To analyze whether the i20S regulates PA200, we made use of primary mouse skin fibroblasts isolated from single i20S subunit KO mice^37^. In these cells we reconstituted the respective i20S subunits with a doxocycline-inducible lentiviral expression system (Figure 7C). Analysis of PA200 RNA expression in i20S single KO and reconstituted cells (virus+ Dox), however, did not reveal any regulation of PSME4 by single i20S subunits (Figure 7C). Our results thus suggest that PA200 has the potential to regulate the relative amounts of intermediate 20S complexes by a yet unknown mechanism while the i20S single subunits do not affect PA200 expression.

**Figure 7:**
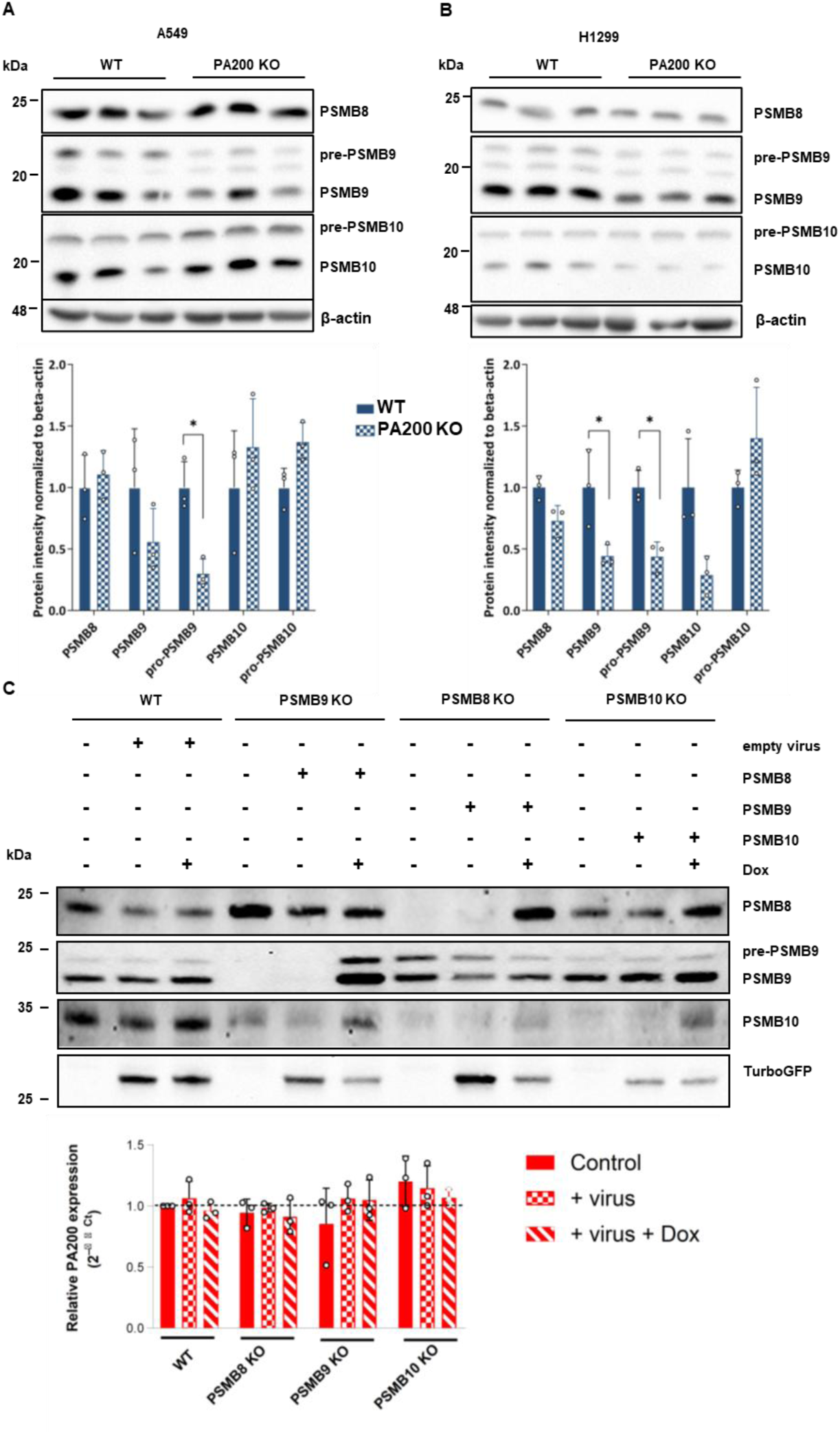
PA200 depletion reduces immunoproteasome expression but immunoproteasome deficiency does not affect PA200 expression. Expression of PA200 and i20S subunits in the lung cancer cell lines A549 **(A)** and H1299 **(B)** upon CrispRCas9-mediated depletion of PA200. Three different WT and PA200 KO clones were analyzed for expression of PA200 and PSMB8-10 as well as for the proteasomal α subunits 1-7. Blots were normalized to β-actin and densitometric analysis shows quantification of expression relative to average WT levels set to 1 (one-sample t-test, n=3). **(C)** primary mouse skin fibroblasts were prepared from PSMB8, PSMB9, and PSMB10 KO mice and compared to cells isolated from WT mice. Single subunits were reconstituted into the respective KO cells using a Dox-inducible lentivirus expressing the single mouse i20S cDNA together with green-fluorescent protein (GFP). WT cells were infected with an empty GFP-expressing lentivirus. Expression of the i20S subunits was analyzed by immunoblotting for the respective subunits. Detection of GFP served as a control for effective lenti-viral transduction of cells. PA200 (PSME4) RNA expression was analyzed by RTqPCR under each condition and expression was normalized to the house-keeping gene Ribosomal protein L19 (RPL19) and to the WT control using the 2^-ΔΔCT^ method (one-sample t-test, n=3 independent experiments).

## Discussion

The i20S proteasome and PA200 were shown to be co-expressed on a cellular level under pathological conditions including patient-derived non-small-cell lung carcinoma^14,27^ and lung fibrosis^19,39^. In this work, we characterized this poorly characterized interaction of PA200 with the i20S in detail and compared it to s20S-PA200 interactions using structural, proteomic and cell biological approaches.

Of note, the cryo-EM structure of the singly-capped i20S-PA200 complex significantly differs from that of the s20S-PA200. Despite an overall very similar fold and opening of the gate, our structural analysis uncovered several striking differences. Firstly, we observed in our cryo-EM data a general bend of the singly capped i20S-PA200, not seen in the three recent s20S-PA200 structures^11,12,23^ and also never observed in any other proteasome activator bound 20S complexes ^23,24,40^. Second, this bend is accompanied by an increase of up to 3.5 Å in the outer diameter of the unbound side of the i20S compared to the s20S, displacing atoms up to 5.4 Å. Using mass photometry, we obtained evidence that the abundance of the singly- and doubly-capped complexes relative to the total proteasome particles was higher for the i20S compared to the s20S. Our data thus suggest that this allosteric bending increases the occupancy of the second PA200 molecule. Such preferred formation of doubly-capped PA200-i20S complexes would thereby indirectly shift the composition of mixed proteasome complexes in the cell. Mixed proteasome complexes consist of the 20S catalytic core with two different proteasome activators attached^2^. This might have implications for cellular function. Third, we detected differential allosteric effects on the catalytic active sites upon binding of PA200 to i20S. These structural shifts were associated with an increased activation of the i20S compared to the s20S upon PA200 binding, as shown by *in vitro* activity assays. More precisely, we observed that PA200 induced a four to ten-fold increase in trypsin- and chymotrypsin-like activities but also a significant activation of the caspase-like activity, although this activity is moderate in the i20S. The activation effect of PA200 on s20S is also significant for the three types of activities but two to five times lower, depending on the substrate peptides used. These divergent data for PA200-mediated activation of i20S and s20S might partly explain the conflicting reports of previous cellular studies on the activation of all three proteolytic activities by PA200 (discussed in detail in^10^). Lastly, we detected the presence of two phytic acid molecules in i20s-PA200 complexes whereas s20S-PA200 complexes associated with both IP6 and 5,6[PP]2-IP4. We show here that IP6 further enhances the activation of the i20S by PA200 to an even larger extent than the s20S, suggesting that it probably acts as a cofactor. Importantly, the trypsin-like activity is most prominently activated by IP6 for both 20S subtypes but to a larger extent for the i20S-PA200 complexes that harbour two IP6 molecules. The binding of IP6 to the positively charged substrate entry channels in PA200 likely promotes the recognition and/or entry of basic substrates potentially altering substrate specificity.

For PA200 to preferentially bind and activate the i20S, it needs to be coexpressed in the same cell and cellular compartment, i.e. the nucleus. We confirmed enriched assembly of i20S-PA200 over s20S-PA200 complexes in testis - an organ that highly expresses PA200 as well as the i20S in about one fourth of all 20S complexes^29^. Coexpression of PA200 and the i20S subunits, however, seems to be restricted depending on the tissues and cells, as determined in human proteome map and human cell atlas data. Moreover, we showed that cellular conditions that upregulated PA200 (e.g. the profibrotic cytokine TGF-β1) or the i20S (e.g. IFNγ, bacterial infection) did not result in concerted transcriptional regulation, suggesting that the proteasome activator PA200 and the catalytic subunits of the i20S are differentially regulated on the transcriptional level. Several studies also observed downregulation of PA200 upon bacterial or viral infection as summarized in^10^. This raises the intriguing possibility that i20S function is fine-tuned by differential expression of proteasomal activators. Regulation of i20S function has been demonstrated for the PA28αβ activator, which is co-regulated with the i20S catalytic subunits by interferons and which is also constitutively co-expressed with the i20S in immune cells^38,41–43^. This activator has been shown to preferentially bind to the i20S^9^ and activate the proteolytic activities of the i20S to foster generation of MHC-I antigenic peptides that activate CD8 T cell responses, e.g. upon viral infections^44–46^. The other member of the PA28 family of activators, i.e. PA28γ, is also able to bind and activate the i20S^9,47^. PA28γ, however, is exclusively expressed in the nucleus and thereby only binds to nuclear i20S. PA28γ has been shown to destroy antigenic peptides which are derived from nuclear pioneer translation products and that serve as an important source for tumor-derived antigenic peptides^48^. As PA28γ is upregulated in many types of tumors it may thereby help tumors to escape from immune surveillance^48^. The lab of Yifat Merbl has recently suggested a similar function for PA200^14^. This study showed reduced inflammatory activation of the i20S by addition of recombinant PA200 to cell extracts. At the same time, PA200 bound strongly to cytokine-induced i20S in pulldown experiments, which is in line with our data. Moreover, silencing of PA200 in the lung cancer cell line A549 reduced the diversity of the antigenic peptides pointing towards an inhibitory function of PA200 in MHC-I antigen presentation^14^. This assumption, however, is not supported by our structural, mass photometry and *in vitro* activity data which clearly indicate favored binding of PA200 to the i20S resulting in prominent activation of all active sites compared to the s20S. While an increase of the CT-L and T-L activities would favor generation of MHC-I antigenic peptides, activation of the intrinsically low C-L activity in the i20S is detrimental for the production of proper MHC-I ligands and efficient antigen presentation^14,49^. Thus, the ability of PA200 to significantly increase the C-L activity of the i20S could at least partly explain the reduced diversity of presented antigenic peptides and the lack of response to immunotherapy observed in lung carcinoma^14^. Another potential explanation for the discrepancy of the data between our and the Javitt study could be the effect of PA200 on β1i expression. We observed reduced RNA and protein expression of PSMB9 encoding β1i using both genetic depletion as well as transient silencing of PA200 in A549 cells. Hence, our data uncover a novel link between PA200 regulation and cellular composition of the proteasome that has not been reported before. This transcriptional regulation may have major implications for antigenic repertoire generation. As such, reduced expression of β1i in PA200 deficient cells might contribute to an altered MHC-I antigenic diversity as reported by Javitt et al.^14^. Indeed, the replacement of a constitutive subunit by its immunosubunit counterpart, and vice-versa, can result both in the generation or destruction of specific antigenic epitopes^36,50^. For example, the increased generation and presentation of the male Y-chromosome-derived UTY 246-254 epitope is favored by β1i, independently of its proteolytic activity, because it impedes the incorporation of the β1 subunit that is responsible for the destruction of this epitope^50^. A similar transcriptional regulation of i20S catalytic subunit expression has been demonstrated for PA28γ, which has been shown to degrade activated Stat3 thereby counteracting interferon-induced transcriptional activation of β1i and β5i subunits and MHC-I antigen presentation^51^. To conclude, our results reporting a function of PA200 in regulating the i20S subunits expression constitute another explanation for the reported role of PA200 in modulating anti-tumor immunity^14^.

## Supplementary Figures

**Figure S1.**
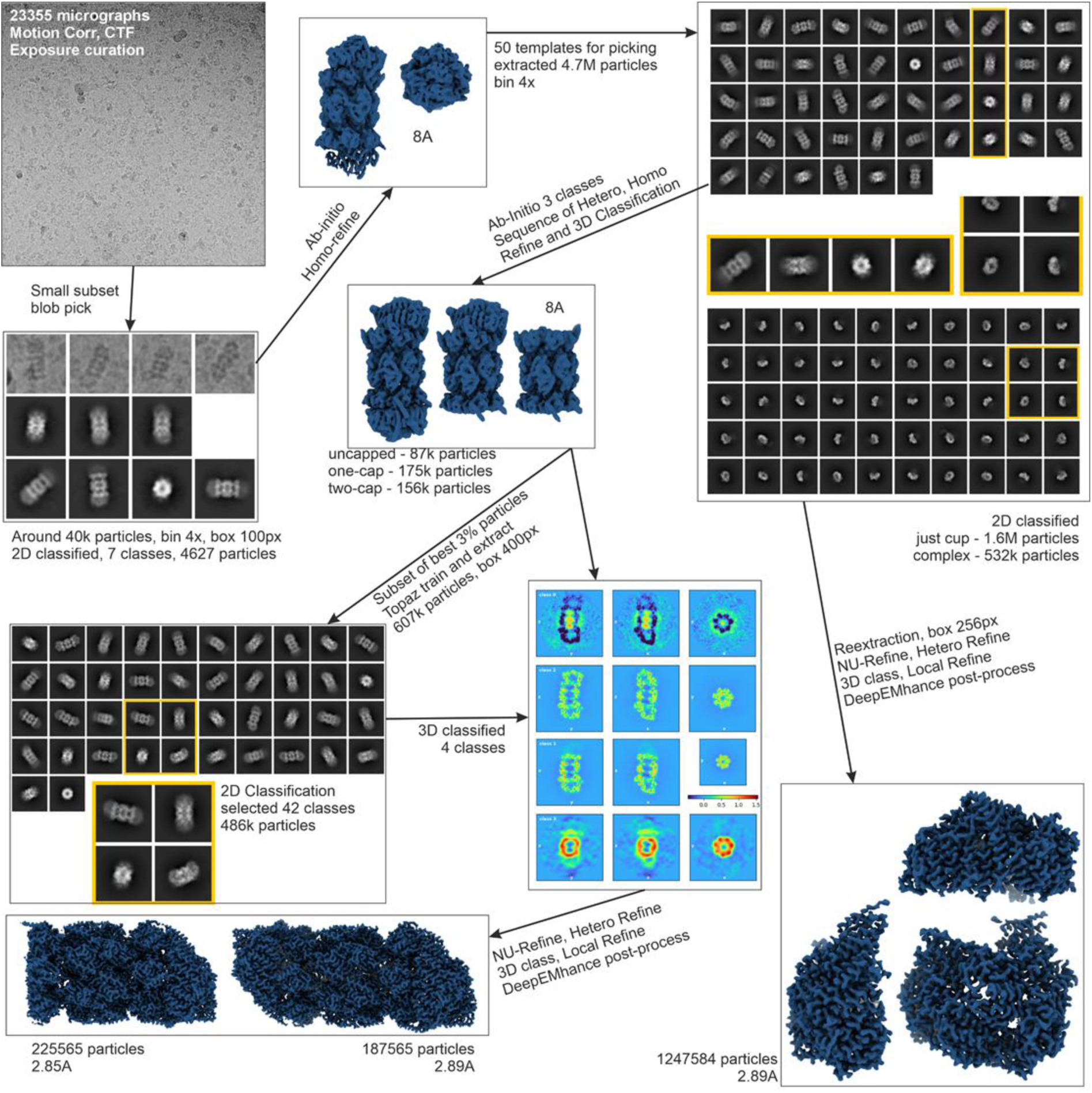
Cryo-EM data processing and 3D classification pipeline.

**Figure S2.**
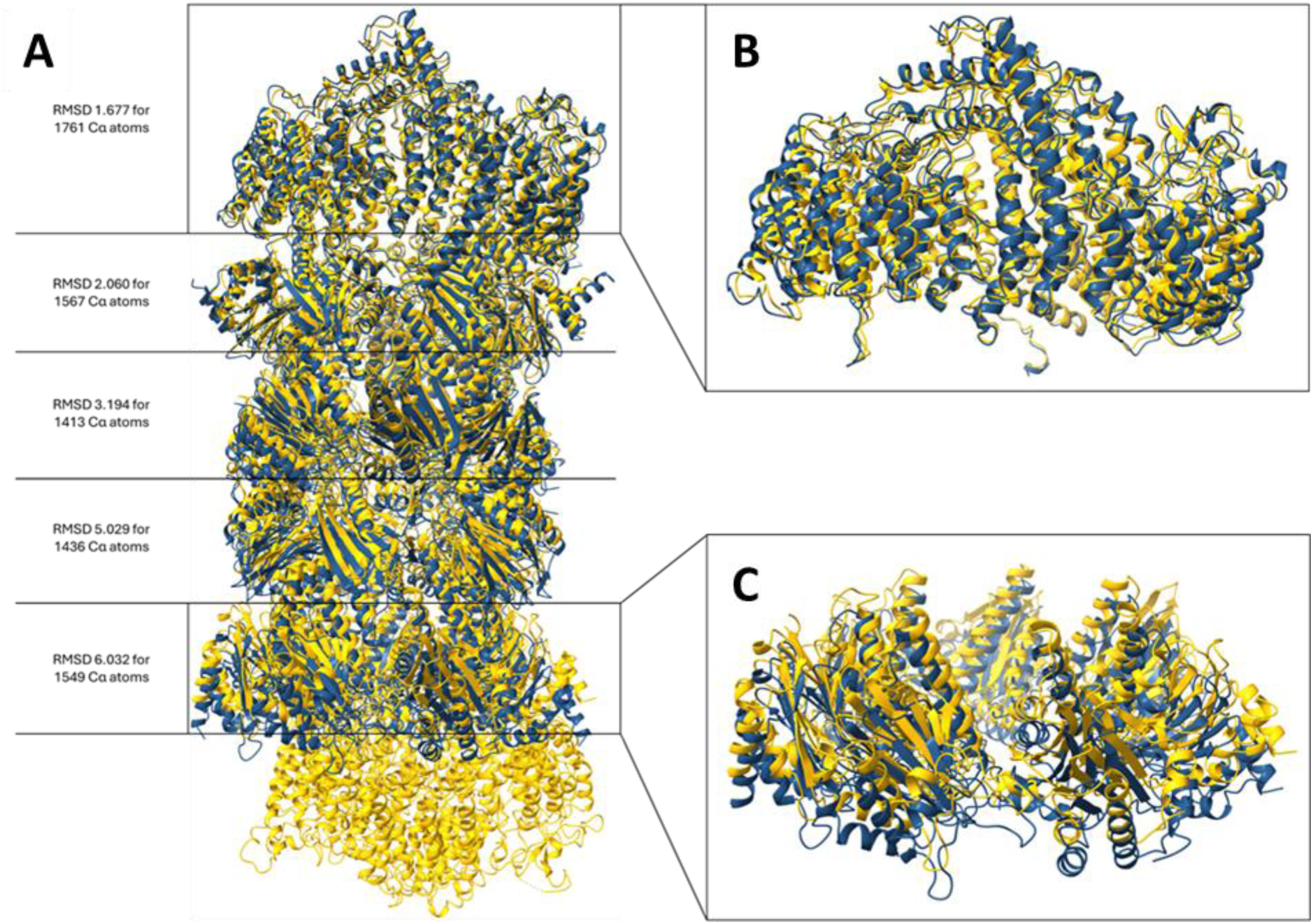
(**A**) Structure of the singly-capped i20S-PA200 complex (in blue) overlaid onto the doubly-capped s20S-PA200 complex (in yellow). The misalignment of the domains far away from the aligned PA200 molecule (**B**) is evident in (**C**).

**Figure S3.**
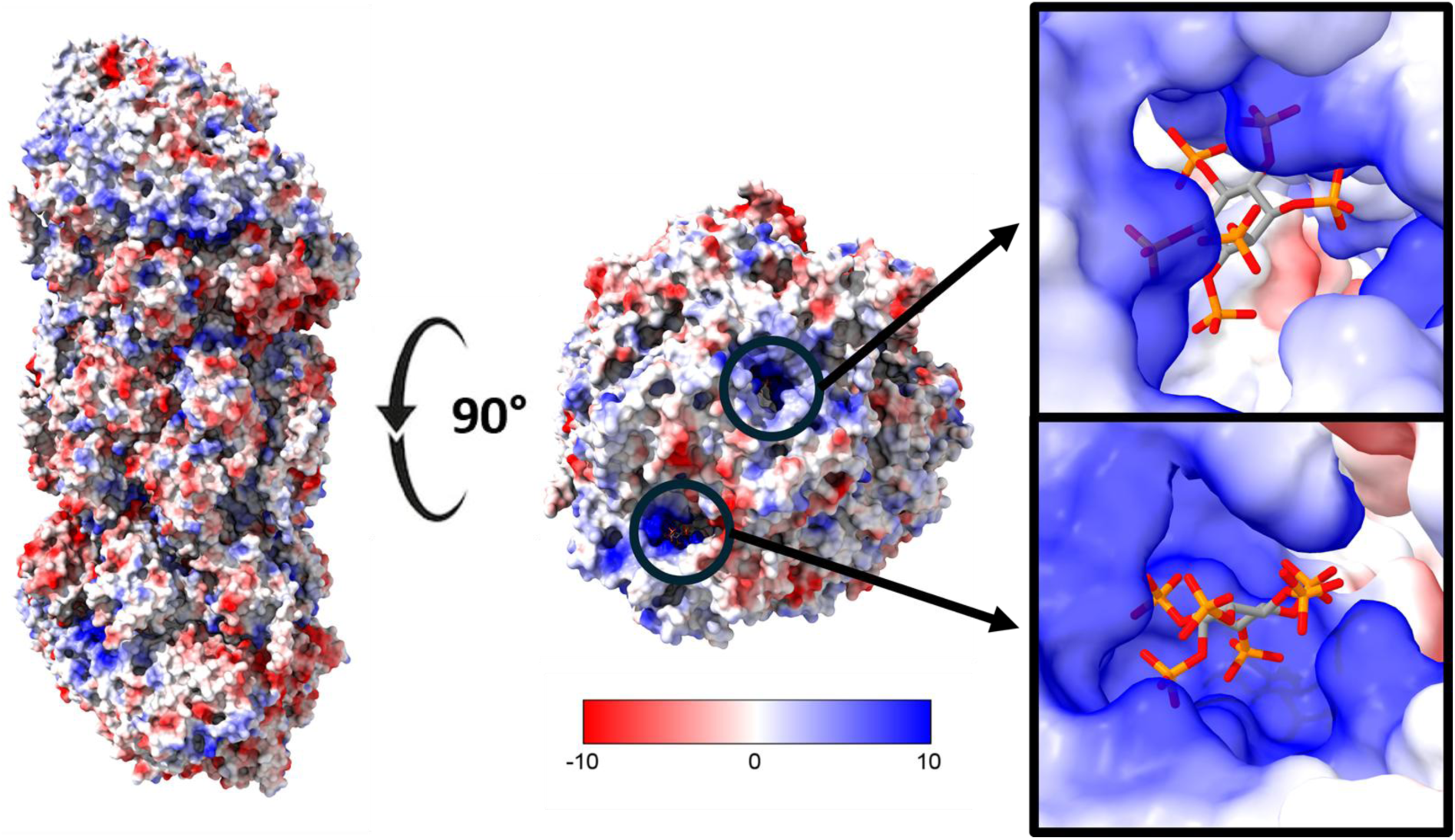
Phytic acid binds to the i20S-PA200 complex. Front and top views of the singly-capped i20S-PA200 complex showing the binding of two IP6 molecules in the positively charged grooves.

**Figure S4:**
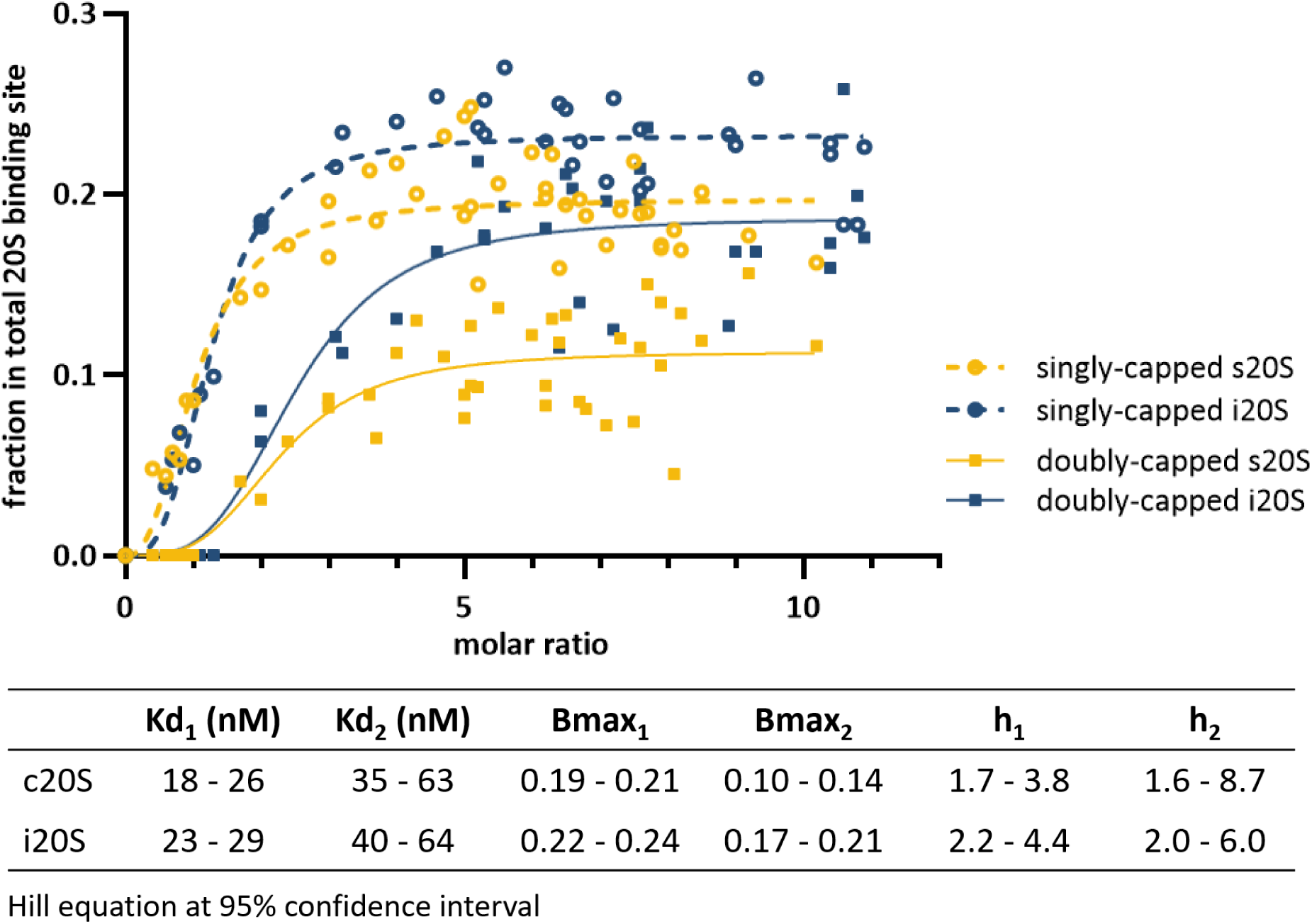
i20S-PA200 doubly-capped particles are preferentially formed compared to s20S-PA200 complexes. Titrations of PA200 to either s20S and i20S showed that the higher occupancy of the i20S is mainly due to a higher population of doubly-capped complexes.

**Figure S5:**
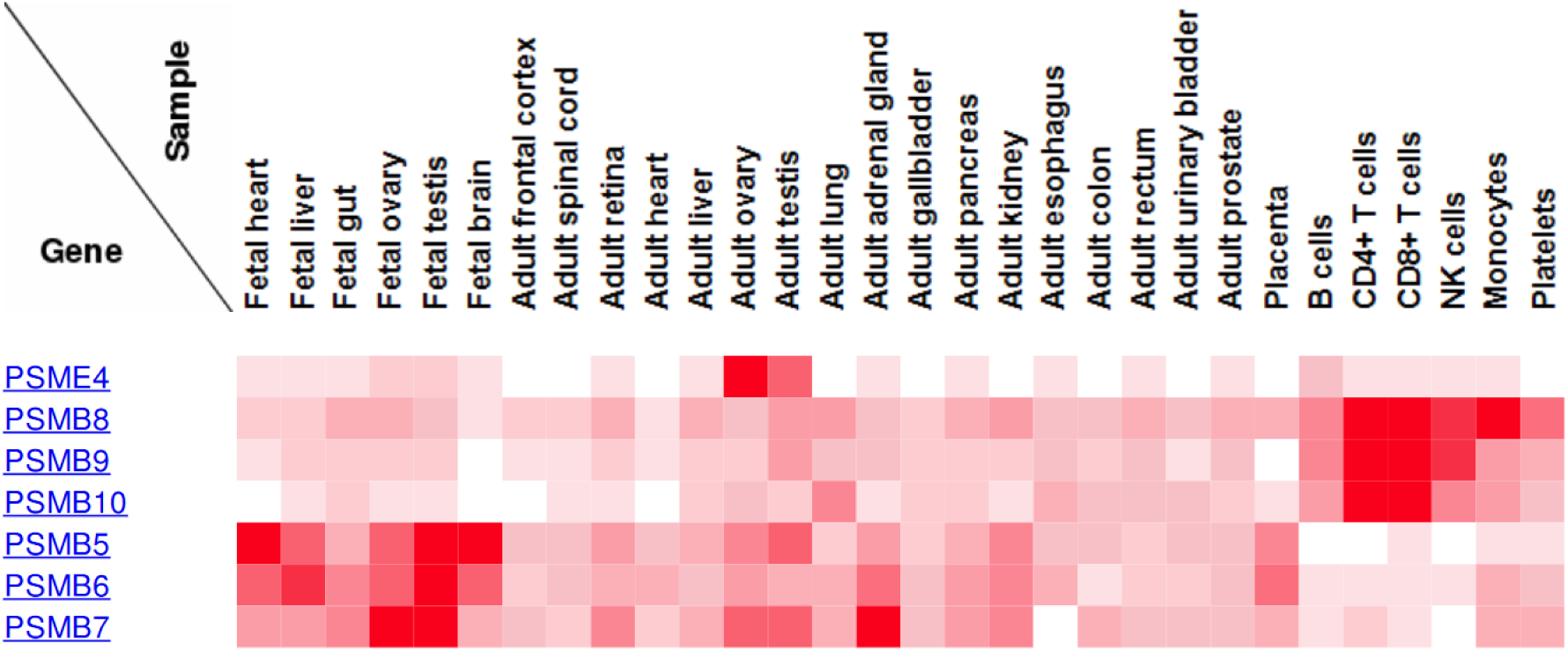
Mapping of the protein expression of PA200 (PSME4) and the catalytic subunits of the i20S (β1i, β2i and β5i, i.e. PSMB8-10, respectively) and s20S (β1, β2 and β5, i.e. PSMB5-7, respectively). Data was retrieved from^33^.

**Figure S6:**
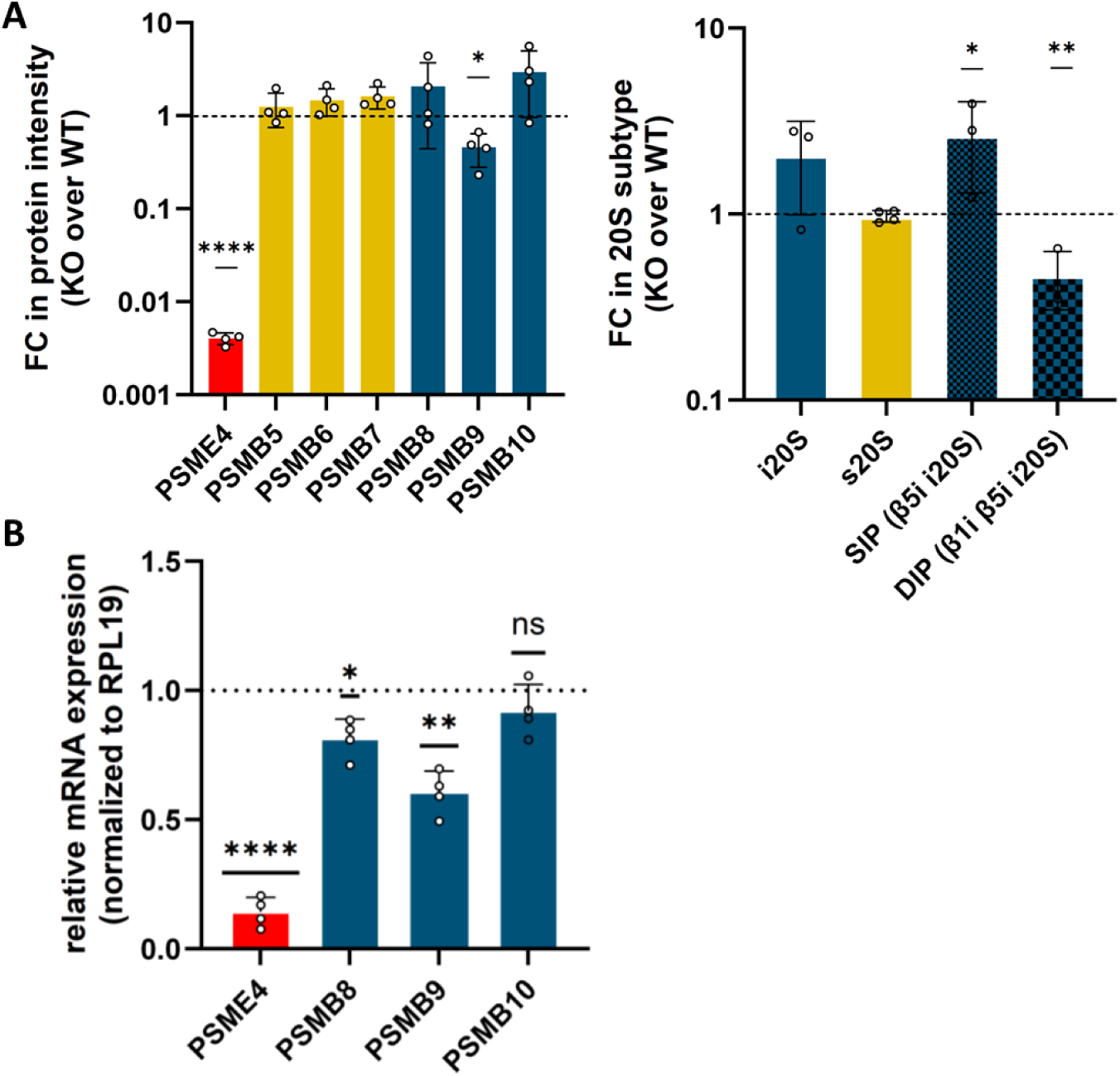
Depletion of PA200 reduces i20S expression and changes the composition of 20S subtypes in lung cancer cells. **A)** Two different WT and PA200 KO clones of A549 and H1299 were used for proteasome immunopurification using the MCP21 antibody. The expression of PA200 and catalytic 20S subunits were quantified by LC MS/MS. For each protein, the log10-fold change of the intensities in PA200 KO over WT was calculated. In this data set, the stoichiometry of each 20S subtype was calculated (as in^38^) and for each 20S subtype, the fold change of the stoichiometry in the PA200 KO over WT was calculated. Bars show quantification of expression relative to average WT levels set to 1 (two-sided paired t-test). *: p<0.05, **: p<0.01. **B)** PA200 was depleted in A549 cells by siRNA-mediated silencing for 72 hr. RNA expression of PSME4 (PA200) and the i20S catalytic subunits PSMB8-10 was analyzed by RTqPCR. RPL19 was used as a housekeeping gene, and the bars indicate mRNA levels normalized to control siRNA transfected cells using the 2^-ΔΔct^ formula. (one-sample t-test, n=4, *: p<0.05, **: p<0.01, ****: p<0.0001).

## Materials and Methods

### Purification of s20S and i20S from HEK293-EBNA Cell Lines

The procedure was done as described in^52^. Proteasome purification was done using MCP21 antibody, produced in-house from a hybridoma, grafted onto cyanogen bromide-activated Sepharose beads. HEK293-EBNA cells, expressing either i20S or s20S, were lysed in a pH 7.6 lysis buffer containing 20 mM Tris-HCl, 100 mM NaCl, 10 mM EDTA, 0.25% Triton X-100, and one tablet of protease and phosphatase inhibitors per 50 mL (cOmplete™ ULTRA Tablets Mini EDTA-free and PhosSTOP, Roche, Basel, Switzerland). Lysate was sonicated with a Vibracell sonicator (10 cycles of 30 s on and 1 min off, at 50% active cycle) and the lysate clarified by centrifugation at 16,000× g for 30 min, and the supernatant was filtered through a 0.22 µm membrane. The filtered lysate was incubated overnight with MCP21 antibody-grafted beads. The following day, the beads were washed with equilibration buffer (20 mM Tris-HCl, 1 mM EDTA, 10% glycerol, 100 mM NaCl, pH 7.6) and eluted using the same buffer supplemented with 3 M NaCl. The eluate was concentrated to 0.5 mL using 100 kDa MWCO centrifugal filters and applied to a size exclusion chromatography on a Superose 6 10/300 GL column with TSDG buffer (10 mM Tris-HCl pH 7.0, 1 M KCl, 10 mM NaCl, 5.5 mM MgCl2, 0.1 mM EDTA, 1 mM DTT, 10% glycerol). Proteasome enzymatic activity was assessed, and active fractions (eluted between 10–14 mL) were pooled. Buffer exchange with equilibration buffer and concentration were performed using 100 kDa MWCO centrifugal filters. Glycerol was added to a final concentration of 20% and aliquots were snap-frozen in liquid nitrogen and stored at −80 °C.

### Recombinant human PA200

Purification of recombinantly expressed human PA200 was done, as recently described^27^. The isolated PA200 was stored at a final concentration of 1 mg/ml to be used for cryo-EM analysis and other *in vitro* assays.

### Cryo EM analysis of i20S-PA200 complexes

i20S was obtained commercially from R&D Systems (Catalog #: E-370, 10 microliter, 25 microgram: <u>Human 20S Immunoproteasome Protein, CF E-370-025: R&D Systems</u>) and complexes were formed upon addition of 8x molar excess of PA200 upon overnight incubation. Before plunge freezing, QUANTIFOIL R2/1 copper grids (200 mesh) were cleaned using a glow-discharger (Leica EM ACE 200) at 8 mA for 60 seconds. 3uL of PA200-i20S sample (0.3 g/l) was added on the grids and plunge-frozen using a Vitrobot Mark IV (Thermo Fisher) set to 95% humidity and 4 °C with 2 seconds for wait time, blot force 5 and 1 second for blotting time. Micrographs were acquired at 300 kV using a Titan Krios G4i (Thermo Fisher; HZ Munich, Germany) equipped with a Selectris energy filter and a Falcon4i direct electron detector. A dataset of 23355 micrographs was collected with 0.95 A pixel size and 1.5–2.5 µm under-focus in 60 frames accumulating 60 e/A^2^ total dose.

### Image processing and volume reconstruction

Cryo-EM datasets were processed using Cryo-EM Single Particle Ab-Initio Reconstruction and Classification (CryoSPARC 4.5.3) software^53^. Imported movies were subjected to motion correction, CTF estimation and manual exposure curation. Around 3% of micrographs were discarded because of unsatisfying CTF resolution estimation, astigmatism or frame motion. Next, a small subset of 200 micrographs was used for preliminary particle picking with a blob picker. After picking, nearly 40.000 particles were 4x binned, extracted with a box size of 100 pixels and 2D classified into 50 classes. 7 classes containing images of supposably full complex (the one with two caps, i.e. PA200 complexes) comprising 4627 particles were used in Ab-initio reconstruction and preliminary Homo refine jobs. The obtained volume was then used to generate 50 templates for template picking on the full dataset. Picked particles were inspected and 4768272 were again 4x binned and extracted with the box size of 100 pixels. Next, a couple of rounds of subsequent 2D classifications were made in 4 parallel groups containing similar numbers of particles. As most of the obtained sequential classes contained just the cup particles, only 532792 particles were used in further processing steps. Sequential 3-class Ab-initio, Hetero-refinement, and 3D classification resulted in three subpopulations for uncapped, 1-cap and 2-cap complexes holding 87822, 175565, and 156772 particles respectively. From the groups, preliminary binned volumes were refined and 3% of the best particles giving the highest CTF resolution estimation were used to train the neural network Topaz picker^54^. Trained model picking resulted in 607937 picked particles, after removal of duplicates. After 2D classification, 486195 particles were selected as believed to be images of the complex. The next steps were a couple of rounds of 3D classification and Hetero refinement to separate the particles of two different complexes, i.e. 1-cap with 225565 particles and 2-cap with 187565 particles. Both densities were NU-refined and Hetero-refined on two identical volumes from NU-refine iteratively. This procedure increased the resolution and was terminated when the resolution of NU-refinement was worse than the step before. The best resolution density was again locally refined and post-processed with DeepEMhancer^55^. Finally, 2.85Å and 2.89Å volumes were obtained for 1-cap and 2-cap complex, respectively.

### Model building and refinement

The PA200 model from PDB 6REY was extracted and fitted into electron density map using Dock in map tool in PHENIX suite^56^. The backbone geometries and orientation of the side-chains were then refined and model was manually rebuilt using Coot^57^. We iteratively corrected steric clashes, Ramachandran and rotamer outliers manually in Coot followed by further refinement using phenix.real_space_refine. Detailed model evaluation was done using Molprobity^58^.

### Proteasome activity assays

The proteasome substrate-based activity assays were performed in a 384-well black plate (Greiner Bio-One, UK). Purified 20S proteasome sample at 0.28 μM (either s20S or i20S) was mixed with 0.56 μM recombinant PA200 regulator in different ratios and left to interact for 30 min at room temperature (22°C). After they were allowed to interact, the complexes were diluted with 50 mM Tris HCl pH 7.6 to a concentration of 3 nM 20S. As a control, both s20S and i20S, was diluted to a concentration of 3 nM. All diluted samples were distributed into wells in aliquots of 20 μL, to which 20 μL of 400 μM fluorogenic peptide substrate was added (prepared in 50 mM Tris-HCl pH7.6): Suc-LLVY-AMC, Boc-LRR-AMC or z-LLE-AMC, to probe for chymotrypsin-, trypsin- or caspase-like activities, respectively. The final proteasome concentration in the reaction was 1.5 nM. The kinetic assays were performed at 37 °C in a CLARIOstar Plus spectrofluorometer (BMG Labtech) over 60 min with one reading every 5 min, at 360 nm for excitation and 460 nm for emission. The slope of the kinetic assay (increase in fluorescence intensity over time) was used to measure proteasome activity. Only slopes at their steepest incline were considered (saturating conditions), after 15-minute temperature equilibration time. At least 5 timepoints were used for slope calculations.

For analysis of the effects of phytic acid on proteasome function, 0.5 μM PA200 was incubated with 1.5 mM Inositol hexakisphosphate (or IP6, Sigma-Aldrich ref P8810) at room temperature for an hour. i20S (purified from HEK-EBNA cells) was then incubated with PA200 (with and without IP6) at 28 nM and 400 nM, respectively, at room temperature for another hour. The complexes were diluted with 50 mM Tris HCl pH 7.6 to a concentration of 2 nM i20S and assayed for activity in 384-well black plates, as described above using the following substrates: Suc-LLVY-AMC (Bachem), Ac-ANW-AMC (UBPBio), Ac-PAL-AMC (UBPBio), Boc-LRR-AMC (Enzo), z-LLE-AMC (UBPBio), Ac-nLPnLD-AMC (Biosynth), or Ac-GPLD-AMC (Biosynth) to probe for chymotrypsin-, β5i-, β1i-, trypsin- or caspase-like activities, respectively.

### Mass photometry

Proteasome (i20S purified from HEK-EBNA, as described in Fabre et al^952^ or s20S purchased from Enzo) was titrated with PA200 for mass photometry as follows: Tris 50 mM pH 7.6 was added to the empty tubes on ice, to compensate for different volumes cause by different theoretical protein ratios (1:0 to 1:12). 20S proteasomes were diluted to 0.2 g/L (0.28 µM) and added to each tube. Recombinantly produced human PA200 was then added to the tubes to form required ratios. Tubes were incubated for 2 h on ice. The complexes were diluted six times immediately before measurement on the Mass photometer (Refeyn Ltd). Acquisition was done using a large field of view, with 60 s recording and default video settings. The acquisition was calibrated using a standard mix of IgG (150 kDa) and thyroglobulin (660 kDa) in Tris-HCl buffer. The data was analyzed in DiscoverMP 1.2 using default settings. The histograms were manually fitted with Gaussian distribution and the counts for each population (PA200 approximately at 200 kDa, PA200 dimer at 400 kDa, 20S at 700 kDa, 20S-PA200 at 900kDa, and PA200-20S-PA200 at 1100 kDa) were recorded.

Tables of counts vs nominal molecular weight were then processed in Excel (Microsoft Office) and graph visualisation and affinity calculation was done in GraphPad Prism.

The experimental molar ratios were obtained based on the actual counts. The fractions of singly- and doubly-capped 20S in total binding site were calculated by obtaining the counts for the singly-capped 20S or 2 x doubly-capped 20S divided by the total possible 20S binding site, which was twice (i.e. two binding site per 20S) the total 20S population (i.e. 20S alone, singly-, and doubly-capped 20S). These two fractions were then summed up to obtain the total 20S occupied binding site. With the fractions of singly- or doubly-capped 20S vs. the experimental molar ratio curves, non-linear fitting was done using Hill equation in GraphPad Prism giving the Kds and Bmax values at 95% confidence interval.

### Co-IP of proteasome complexes and proteomics - interactomics

Co-immunoprecipitation was done using anti-PA200 antibodies or anti-α2 proteasome subunit antibodies for hPLF samples and for testis samples. The proteasome complex enrichment procedure for cultured cells is described in detail in ^29^.

Briefly, the material is homogenized using a Potter device (testis) or a cell scraper (hPLFs) in native buffer, then centrifuged to remove cell debris. Magnetic Protein G beads were incubated with anti-PA200 or anti-α2 proteasome subunit antibodies and the antibodies were then cross-linked to the beads via short 0.1% formaldehyde treatment. Clarified lysate was then incubated with the magnetic beads coated in antibodies overnight at 4°C. The beads were washed 4 times in lysis buffer, and enriched proteasome complexes are eluted from the beads using SDS. The samples were prepared for proteomics using S-Trap protocol, as described by the supplier (Protifi).

LC-MS method with bioinformatic approach for interactomics is also described in^29^. Briefly, a standard nano-flow reverse phase LC-MS proteomic method was used, in DDA mode, on an Ultimate 3000 LC system and an Orbitrap Fusion MS instrument. Bottom-up LC-MSMS and Bottom-Up bioinformatic searches were done as described in the Appendix 1 of Zivkovic et al. 2022^29^, with the exception that the fasta-derived *in silico* library was human, from the Uniprot Proteome database. The proteomic search was done using the Proline suite, an easy-to-use, open source bioinformatic tool developed in-house^59^.

For the analysis of the PA200 KO cell cultures, same lysis conditions, co-immunoprecipitation and protein digestion procedure was followed as described in detail in^29^, however, a different instrumental setup was used for the proteomic analysis.

Digested peptide extracts were desalted on an Evotip C18 EV2001 tips (Evosep) and were analyzed by online nanoLC using an UltiMate 3000 RS nano-LC system (Thermo Scientific) coupled with a TimsTOF SCP mass spectrometer (Bruker). Peptides were separated on a C18 Aurora column (25 cm x 75 µm ID, IonOpticks) using a gradient ramping from 2% to 20% of B in 30 min, then to 37% of B in 3 min and to 85% of B in 2 min (solvent A: 0.1% Formic Acid (FA) in H2O; solvent B: 0.1% FA in Acetonitrile (ACN)), with a flow rate of 150 nL/min. MS acquisition was performed in DIA-PASEF mode on the precursor mass range [400-1,000] m/z and ion mobility 1/K0 [0.64-1.37]. The acquisition scheme was composed of 8 consecutive TIMS ramps using an accumulation time of 100 ms, with 3 MS/MS acquisition windows of 25 Th each. The resulting cycle time was 0.96 s. The collision energy was ramped linearly as a function of the ion mobility from 59 eV at 1/K0=1.6 Vs.cm−2 to 20 eV at 1/K0=0.6Vs.cm−2. The raw data was searched and quantified with DIA-NN 1.9, using a predicted library from UniProt human reference proteome. The result files were then imported into Proline^59^ for validation and label-free quantitation with a False Discovery Rate (FDR) of ≤ 1.0 %, peptide length range set at 7-30 and precursor charge states of 2+ and 3+. For statistics, a t-test p-value threshold of 0.01 was applied at both peptide and protein levels. Only specific peptides were used for quantification, and the median ratio fitting was chosen as the abundance summarizer method. Normalization, missing values inference (if strictly less than three values, 5% centile was applied), t-test and z-test (p-value Benjamini-Hochberg correction) were also applied.

### Cell isolation, culture and treatment

A549 and H1299 non-small cell lung cancer cell lines were grown at 37 °C and 5% CO_2_ in a humidified incubator in DMEM/F12 or RPMI-1640 medium, respectively, with supplementation of 10% FBS and 100 U/mL penicillin/streptomycin. CRISPRCas9-mediated genomic depletion of PSME4 (gene name for PA200) has been described previously by us in detail^27,59^.

Primary human lung fibroblasts (phLF) obtained from organ donors were used as described previously^19^. Cells were cultured in MCDB medium supplemented with 10% (v/v) FBS (Biochrome), 100 U/mL penicillin/streptomycin (Gibco, Thermo Fisher Scientific), 2 mM L-glutamine (Thermo Fisher Scientific), 5 µg/mL insulin (Thermo Fisher Scientific), 2 ng/mL basic-FGF (Thermo Fisher Scientific), and 0.5 ng/mL human EGF (Sigma-Aldrich). Cells were allowed to grow to 90% confluency before splitting. For subculturing, cells were rinsed with PBS, treated with 0.25% Trypsin-EDTA (Sigma) for 4–5 minutes at 37 °C, re-suspended in fresh culture medium, and transferred to new dishes. All experiments were conducted using phLF between passages 3 and 5. Prior to treatment of phLF, 0.3 x 10^6^ cells were seeded into 10 cm plates for 24 h. The cells were then synchronized by culturing them in modified MCDB medium (see above) containing 1% FBS for 24 h. Following synchronization, the medium was refreshed, 5 ng/ml TGF-β1 or 75 U/ml IFNγ were added into the culture medium for 48 h. Cell harvest was achieved by washing the cells with PBS and then detaching the cells by adding 0.25% Trypsin-EDTA and incubating the plates for 5 minutes at 37°C. The process was stopped by adding culture media with 10% FBS, and the collected cells were then centrifuged in a 15 ml falcon tube at 5000 rpm for 5 minutes. Those cell pellets were washed once more with PBS in 1.5 ml Eppendorf tubes before their storage at -80° until further use.

The method used for *primary mouse skin fibroblast* isolation was reported previously (doi: 10.3791/2033) with minor modifications regarding the culture medium where DMEM was supplied with 10% FBS (Capricorn) and the antibiotic 1×Antimycotic (Gibco, 15240062) was used.

Bone-marrow derived mouse macrophages (BMDM) were generated from C3HeB/FeJ mice as previously described^60^. BMDMs then were cultured at 5 x 10^5^ cells/well and incubated overnight in 24-well plates (Nunc-surface). Subsequently cells were infected with *Mycobacterium tuberculosis* H37Rv at a multiplicity of infection (MOI) of 3:1 for 24 h. Total RNA of cells lysed in Trizol (peqGOLD TriFast, VWR International) was extracted by use of DirectZol RNA MiniPrep (Zymo Research) according to the manufacturer’s instructions. For reverse transcription of RNA, the Maxima First Strand cDNA Synthesis Kit for RT-qPCR (Thermo Fisher Scientific) was used.

### RNA isolation of cells

Total RNA from cells was extracted using phenol-chloroform extraction with TRIzol Reagent invitrogen). Cells were resuspended in TRIzol Reagent, followed by vigorous mixing with chloroform in Phasemaker tubes. The samples were then incubated for 5 minutes and centrifuged at 16,000 x g for 5 min at 4 °C to achieve phase separation. The upper aqueous phase was carefully transferred to a new tube, and 500 µL of isopropanol was added. The samples were incubated for 10 minutes on ice. RNA was pelleted by centrifugation at 16,000 x g for 10 min at 4 °C. The supernatant was discarded, and the RNA pellets were washed with 70% (v/v) ethanol. After drying on ice, the RNA pellet was dissolved in 30 µL of nuclease-free water and incubated in a heat block at 55°C for 10 min, and the RNA concentration was measured at 260 nm using the NanoDrop 1000 (ThermoFisher).

### Reverse transcription (RT) of mRNA and RT-qPCR

For reverse transcription, 0.5 to 1 µg RNA was diluted to 14 µL with nuclease-free water and combined with a 6 µL mastermix of Maxima First Strand cDNA Synthesis Kit for RT-qPCR according to the manufacturer’s instructions (ThermoFisher). The reaction was terminated by heating at 85°C for 5 min. The samples were then diluted 1:5 with nuclease-free water.

Quantitative real-time PCR was carried out using a SYBR Green LC480 system (Roche). Each well in the 96-well plate contained a mixture of 2.5 µL cDNA and 5 µL LC480 SYBR Green I Master mix (Roche), along with 2.5 µL of forward and reverse primers, resulting in a final concentration of 0.5 µM. All samples were measured in duplicate, and plates were centrifuged at 1000 rpm for 2 minutes before beginning the measurement, following the standard protocol of the Light Cycler 480II (Roche). Gene expression levels were normalized to the housekeeping gene hypoxanthine-guanine phosphoribosyltransferase (HPRT). Relative gene expression was calculated using the ΔΔCT method.

### Primer sequence table

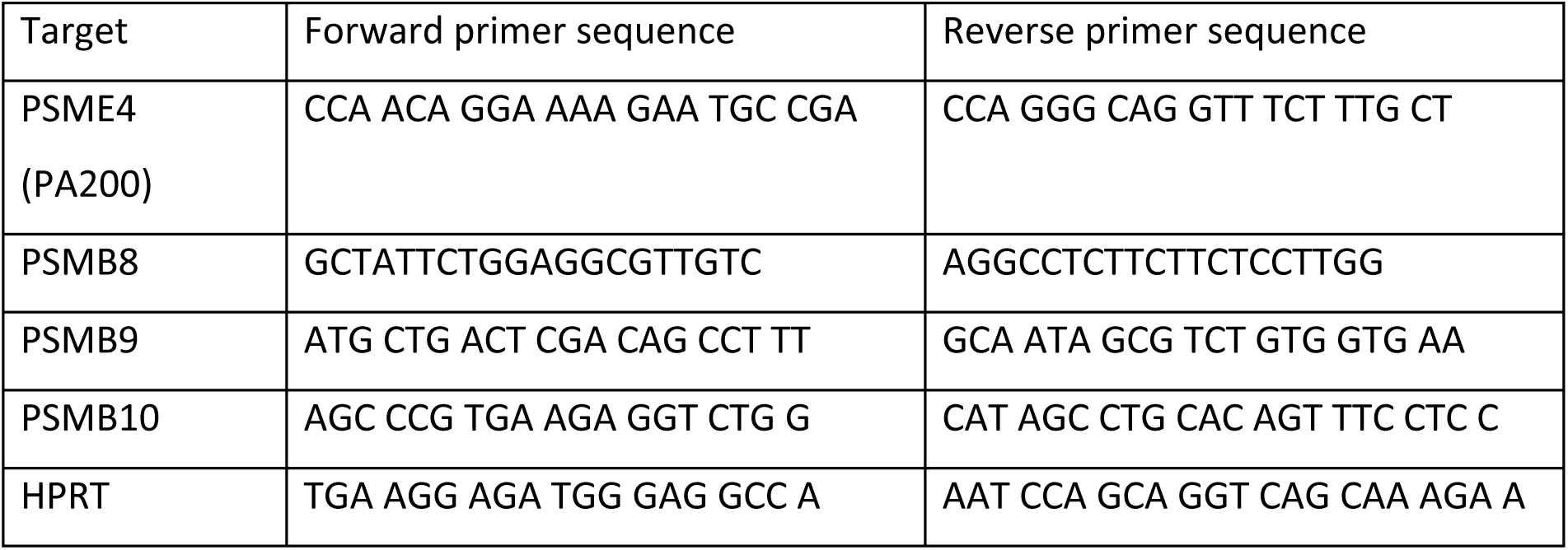

### Protein extraction and quantification followed by SDS-PAGE and Western blot analysis

In order to evaluate intact and active proteasome complexes, cell pellets were lysed under non-denaturing conditions. Cell pellets were resuspended in OK40 buffer (50 mM Tris-HCl, 5 mM MgCl_2_ 10% Glycerol, 0.05% NP-40, 2mM ATP) containing a 1x cOmplete™ protease inhibitor cocktail (Roche) and 1x phosphoSTOP (Roche) and lysed through vigorous pipetting followed by a 20 min incubation on ice with additional vigorous pipetting and vortexing. The lysates were then centrifuged at 15,000 rpm for 20 min at 4 °C. The supernatant was transferred to new tubes for immediate protein concentration determination using a BCA assay. A bovine serum albumin (BSA) calibration curve, with concentrations ranging from 0 to 2 µg/µL in PBS, was used as a standard for protein quantification. To perform the assay, 20 µL of BSA standard, 2 µL of protein lysate or pure lysis buffer diluted 18 µL in PBS were combined with 200 µL of BCA reagent, following the manufacturer’s instructions (Thermo Fisher Scientific). After a 30-minute incubation at 37 °C, absorbance was measured at 562 nm using a plate reader for subsequent protein concentration calculation.

15 µg of protein were used per sample for western blotting, and 25 µg for ABP assay. Protein extracts were diluted to equal volumes with Milli-Q® water, then mixed with 6x Laemmli sample buffer. The protein mixture was heated to 95 °C for 10 min to denature the proteins. For protein electrophoresis, samples were loaded onto 12% SDS polyacrylamide gels. Protein samples, along with a Protein Marker (#26616, ThermoFisher) were loaded onto SDS polyacrylamide gels. Electrophoresis was carried out using Bio-Rad gel running chambers. Following electrophoresis, proteins were transferred to a polyvinylidene fluoride (PVDF) membrane (Bio-Rad) using the tank immunoblotting method. The membrane was first activated in pure methanol, and the transfer was carried out at a constant current of 250 mA for 90 min or an overnight transfer of 40 mA for 960 min. To block nonspecific binding, the PVDF membrane was incubated in Roti®-Block solution (Carl Roth) for 1 h. The membrane was then incubated with the primary antibody, diluted in Roti-Block solution, either overnight at 4 °C or for 1 h at room temperature (RT). Afterwards, the membrane was washed three times with PBST (PBS, 1% Tween-20) for 5 min each and incubated with a horseradish peroxidase-conjugated secondary antibody, diluted 1:20,000 in PBST, for 60 min at RT on a shaker. The membrane was then washed three more times with PBST for approximately 20 min in total, and proteins were detected using a chemiluminescent substrate according to the manufacturer’s instructions. Protein signals were detected with the iBright CL750 (ThermoFisher). The following antibodies were used: anti-LMP2 (ab242061, Abcam), anti-LMP7 (ab3329, Abcam), anti-MECL1 (ab183506, Abcam), anti-TurboGFP (TA150041, OriGene), anti-GAPDH (14C10, Cell Signaling), anti-PA200 (NBP1-22236, Novus Biologicals).

### Lentivirus design and production

Lentiviruses were constructed using the pCW57-MCS1-P2A-MCS2 (GFP) transfer vector (#80924, Addgene). The cDNAs for the three different immunoproteasome subunits LMP7, LMP2 and MECL-1 were amplified from mouse embryonic fibroblasts using primers containing restriction sites for Nhe I-mediated cloning into the vector. The vector pCW57-MCS1-P2A-MCS2 (GFP) was linearized with the restriction enzyme NheI (NEB, #R3131), whose restriction site is located upstream of the P2A region. The cDNAs of the individual immunoproteasome subunits were then integrated into the linearized vector after digestion with NheI using the NEBuilder® HiFi DNA Assembly Master Mix (NEB, #E2621). The construct allows inducible expression of the gene of interest by addition of Doxycycline (Dox). The activity of the tetracycline response element (TRE) promoter is inhibited by a reverse Tetracycline repressor (rTetR), which is expressed by a downstream hPGK promoter, when Dox is absent. In contrast, addition of Dox allows the release of the rTetR from the TRE promoter thereby allowing expression of the gene of interest. The turboGFP gene is driven independently of Dox by the hPGK promoter and allows sorting of lentivirus - infected cells.

5×10^6^ 293 HEK-T cells were seeded into 10-cm cell culture plates 24 h prior to the transfection of plasmid. Before plasmid transfection, fresh and pre-warmed DMEM supplied with 10% FBS, 25 mM HEPES and 1% L-glutamine was added to the culture plate. 8 μg of transfer plasmid (pCW57-MCS1-P2A-MCS2 (GFP) was used as control, pCW57-LMP2-P2A-GFP, pCW57-LMP7-P2A-GFP or pCW57-MECL1-P2A-GFP) together with 6 μg pSPAX2 (Addgene) and 4 μg pMD2.G (Addgene) were transfected using PEI (Merck, 919012). Cell culture medium was replaced after 8 h of transfection, and lentivirus-containing medium was collected 48 h after medium replacement. Collected lentivirus-containing medium was filtered with 0.45 μm sterile filters prior to use and stored at -80°C.

### Lentivirus infection and Dox induction

For lentivirus infection, 3×10^5^ WT or immunoproteasome-deficient skin fibroblasts were seeded into 6-cm plates 24 h before infection. On the day of infection, either empty lentivirus or lentivirus containing the cDNAs of β1i (PSMB9), β5i (PSMB8), or β2i (PSMB10) were added to the plates with a multiplicity of infection (MOI) of 3 for 48 h. Simultaneously, polybrene (Merck, TR-1003) with a final concentration of 8 μg/ml was added to the medium to enhance infection efficiency. After 48 h, fibroblasts were then washed with PBS to remove residual polybrene and lentivirus. Afterwards, 1 μg/ml Doxycycline (Dox) (Merck, D5207) was applied to infected cells for 96 h to induce expression of the target gene and turbo GFP. Dox was added to medium every 48 h.

## Data Availability

The mass spectrometry proteomics data concerning A549/H1299 PA200 KO cells and TGF-β treated phLF have been deposited to the ProteomeXchange Consortium via the PRIDE^61^ partner repository with the dataset identifier PXD061729. The anti-PA200 and anti-α2 CoIP proteomics data in testes were previously deposited with the dataset identifier PXD027436^29^.

## Acknowledgements

The study was supported by a BMBF grant to SM and GP (Nr. 16GW0287), a DFG/ANR grant to SM, JB, and MPB (ME2002/6-1, BE1305/9-1, and ANR-PA200_IN_IPF) and by grants from the French National Research Agency (ProFI projects: ANR-10-INBS-08 & ANR-24-INBS-0015), the Région Occitanie, and the REACT-EU program of the European Commision. The work is supported under the Polish Ministry and Higher Education project: “Support for research and development with the use of research infrastructure of the National Synchrotron Radiation Centre SOLARIS” under contract nr 1/SOL/2021/2.

We gratefully acknowledge the provision of human biomaterial from the CPC-M bioArchive and its partners at the Asklepios Biobank Gauting and the Klinikum der Universität München.

## Abbreviations

s20S: standard 20S proteasome
i20S: immunoproteasome (20S)
phLF: primary human lung fibroblasts
GFP: green fluorescent protein
TGF-β1: transforming growth factor β1
IFNγ: interferon-γ
Mtb: Mycobacterium tuberculosis
WT: wildtype
KO: knockout
Dox: doxycycline
CT-L: chymotrypsin-like
T-L: trypsin-like
C-L: caspase-like
RT: room temperature
BSA: bovine serum albumin
RT: Reverse transcription
HPRT: hypoxanthine-guanine phosphoribosyltransferase
RPL19: ribosomal protein L19

